# Neurexins play a crucial role in cerebellar granule cell survival by organizing autocrine machinery for neurotrophins

**DOI:** 10.1101/2020.11.14.383158

**Authors:** Takeshi Uemura, Emi Suzuki, Shiori Kawase, Taiga Kurihara, Misato Yasumura, Tomoyuki Yoshida, Shuya Fukai, Maya Yamazaki, Peng Fei, Manabu Abe, Masahiko Watanabe, Kenji Sakimura, Masayoshi Mishina, Katsuhiko Tabuchi

**Author notes:** Correspondence should be addressed to T.U. or K.T.

## Abstract

Neurexins (NRXNs) are among the key presynaptic cell adhesion molecules that regulate synapse function and formation via trans-synaptic interaction with postsynaptic ligands. Here, we generated cerebellar granule cell (CGC)-specific *Nrxn* triple-knockout (TKO) mice to allow the deletion of all NRXNs. Unexpectedly, most CGCs died in these mice. The requirement of NRNXs for cell survival was reproduced in cultured CGCs. We showed that the axons of cultured *Nrxn* TKO CGCs that were not in contact with the postsynaptic structure had defects in the formation of presynaptic protein cluster and action potential-induced Ca^2+^ influx. Additionally, these cells were impaired in the secretion from axons of depolarization-induced fluorescence-tagged brain-derived neurotrophic factor (BDNF), and the cell-survival defect was rescued by the application of BDNF. Our results suggest that CGC survival is maintained by autocrine neurotrophic factors, and that NRXNs organize the presynaptic protein clusters and the autocrine neurotrophic factor secretory machinery independent of contact with postsynaptic ligands.

## INTRODUCTION

Synapse formation is a multistep process that is triggered after the initial contact of an axon with a postsynaptic cell; however, this process is thought to vary according to the type of synapse^1^. During this process, synaptic cell adhesion molecules called synapse organizers play crucial roles in triggering the formation, organization, and functional specialization of synapses. Neurexins (NRXNs) are among the major presynaptic organizers and interact with various types of postsynaptic cell adhesion molecules including neuroligins (NLGNs), leucine-rich repeat transmembrane neuronal proteins (LRRTMs), and the cerebellin precursor protein 1 (Cbln1)-glutamate receptor δ2 (GluD2) complex^2^. In mammals, NRXNs are encoded by three genes (*Nrxn1, Nrxn2*, and *Nrxn3*). Two alternative promoters in each gene produce a long α-NRXN and a short β-NRXN, and each transcript undergoes alternative splicing at six sites (splice sites 1–6) and at two sites (splice sites 4 and 5), respectively^2^. Although *Nrxn* mRNAs are widely expressed in the brain, the expression patterns of multiple *Nrxn* variants differ according to cell type^3^. The functions of NRXNs in various types of synapse are supported by several physiological evidences. Genetic manipulations of *Nrxn* genes in mice have shown that α-NRNXs and β-NRXNs play different roles in synaptic functions in different types of synapses^4–7^. Triple deletion of *Nrxn1, Nrxn2*, and *Nrxn3* in four different types of synapses, i.e., climbing fiber (CF)–Purkinje cell (PC) synapses, synapses formed by parvalbumin- or somatostatin-positive interneurons on pyramidal layer 5 neurons of the medial prefrontal cortex, and the calyx of Held synapse, yields severe but distinct synaptic phenotypes, ranging from impairments in their distribution and functions to decreases in synapse number^8,9^.

The process of synapse formation between parallel fibers (PFs), which are the axons of cerebellar granule cells (CGCs), and cerebellar PCs has unique characteristics compared with that of other types of synapses, such as those in neocortical and hippocampal pyramidal neurons; i.e., the dendritic spines of cerebellar PCs are formed by cell-intrinsic factors without presynaptic contact^1,10^. Additionally, presynaptic elements of the PFs with vesicular profiles similar to those of synaptic vesicles are also intrinsically formed in the absence of their specific postsynaptic compartments^10,11^. However, the physiological role of these presynaptic-like structures remains unknown, and the cell-intrinsic factors that are necessary for the formation of the presynaptic-like structure have not been identified. Conversely, the synapse organizers that mediate the formation of PF–PC synapses have been identified. Previously, we found that the PF–PC synapses are formed and maintained by a *trans*-synaptic interaction between postsynaptic GluD2 and presynaptic NRXNs through the secreted Cbln1^12^. The functional importance of this ternary complex at the PF–PC synapse has been confirmed by both conventional and conditional *GluD2* or *Cbln1* knockout mice, in which cerebellar long-term depression (LTD) is impaired and approximately half of spines lack synaptic contact with the PF terminal^12–16^. However, in contrast with GluD2 and Cbln1, the physiological roles of the presynaptic-organizing NRXNs in CGCs have not been fully clarified.

In the present study, we generated CGC-specific *Nrxn1, Nrxn2*, and *Nrxn3* triple-knockout (*Nrxn* TKO) mice to clarify the physiological role of NRXNs in CGCs. Contrary to the initial expectation, we found that almost all CGCs died in these mutant mice. This phenotype was entirely different from those of both conventional and conditional *GluD2* or *Cbln1* knockout mice^12–16^. We examined the effect of conditional *Nrxn* TKO in cultured CGCs, and found that NRXNs were required for their survival. The cultured *Nrxn* TKO CGCs had a morphological and functional defect of presynaptic-like structure which is formed independently of postsynaptic contact. Notably, in these cells, the activity-induced axonal neurotrophic factor secretory machinery was severely impaired, and the cell-survival defect was rescued by application of brain-derived neurotrophic factor (BDNF). Thus, our results suggest that NRXNs regulate the functional organization of the presynaptic-like structure and that axonal NRXNs are essential for the self-regulatory machinery of neurotrophic-factor-mediated CGC survival.

## RESULTS

### Generation of CGC-specific *Nrxn* TKO Mice on the pure C57BL/6 genetic background

To investigate the physiological role of NRXNs in PF–PC synapses, we generated CGC-specific *Nrxn* TKO mice. To knock out both *α-Nrxn* and *β-Nrxn* genes, we constructed targeting vectors in which two *loxP* sequences flanked the common last exon of each *α-Nrxn* and *β-Nrxn* gene (Fig. 1a). Using embryonic stem cells derived from the C57BL/6 strain, we generated *Nrxn1^flox/flox^; Nrxn2^flox/flox^; Nrxn3^flox/flox^* mice (Supplementary Figure 1). These mice were crossed with CGC-specific Cre (*GluN2C^+/iCre^*) mice^17^ to yield *GluN2C^+/iCre^; Nrxn1^flox/flox^; Nrxn2^flox/flox^; Nrxn3^flox/flox^* mice, which were termed CGC-specific *Nrxn* TKO mice (Fig. 1b,c). *GkuN2C^+/+^; Nrxn1^flox/flox^; Nrxn2^flox/flox^; Nrxn3^flox/flox^* (floxed *Nrxn*) littermates were used as a control.

**Fig. 1.**
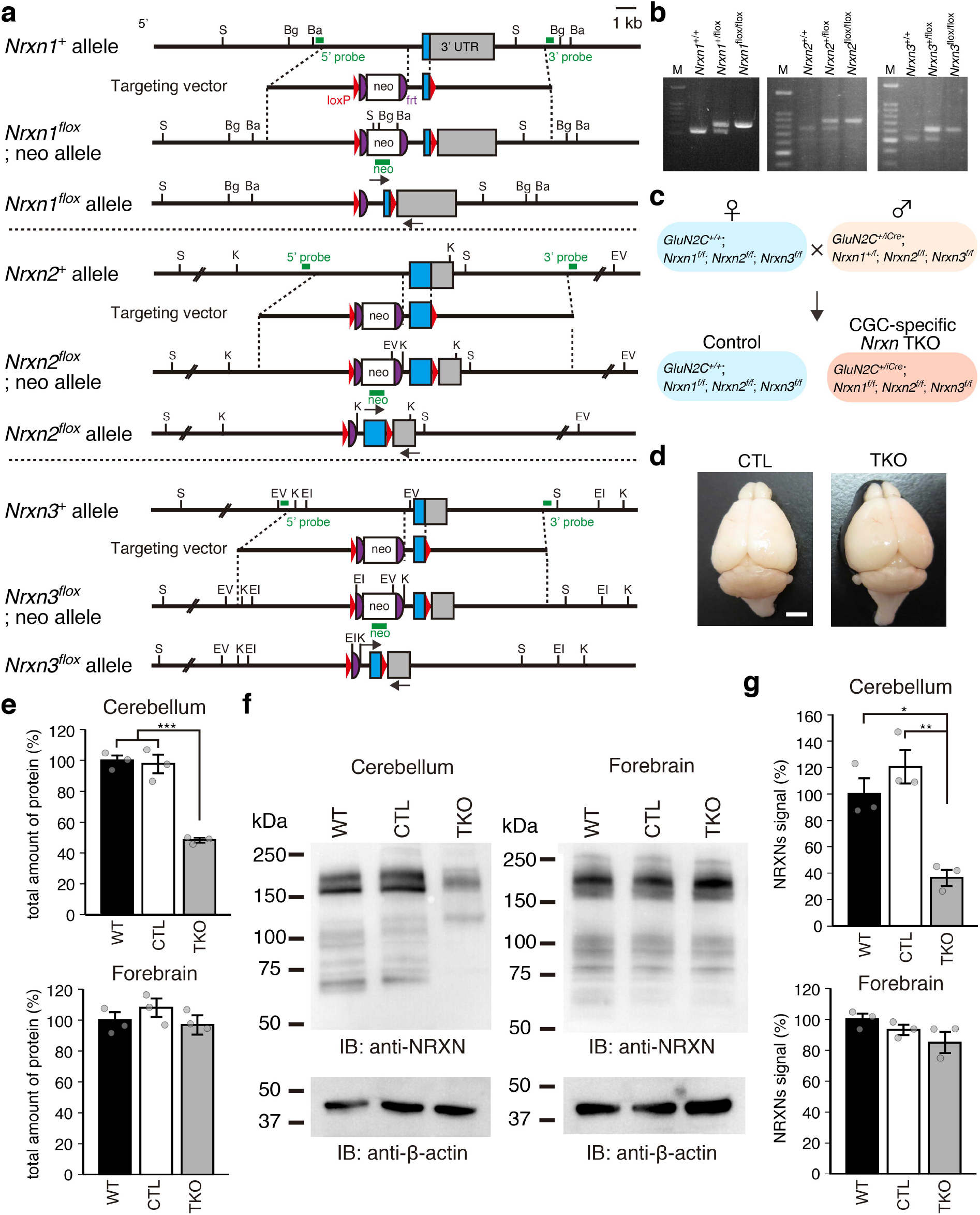
Generation of CGC-specific *Nrxn* TKO mice. **a**, Schema of the wild-type *Nrxn (Nrxn^+^), neo*-inserted alleles (*Nrxn^flox^; neo*), and floxed allele (*Nrxn^flox^*) in each *Nrxn* gene. The arrows indicate the PCR primers in **b**. Ba, *Bam*HI; Bg, *Bgl*II; EI, *Eco*RI; EV, *EcoR*V; K, *Kpn*I; S, *Sca*I. **b**, Agarose gel electrophoresis of the DNA fragments amplified by PCR from *Nrxn^+/+^, Nrxn^+/flox^*, and *Nrxn^flox/flox^* mice. **c**, Schema for the generation of CGC-specific *Nrxn* TKO mice. *Nrxn^flox^* mice were crossed with CGC-specific Cre mice (*Grin2C^iCre^* mice). **d**, Dorsal view of the brain of 8-week-old control (*Grin2C^+/+^; Nrxn1^flox/flox^; Nrxn2^flox/flox^; Nrxn3^flox/flox^*) (left panel) and CGC-specific *Nrxn* TKO (*Grin2C^+/iCre^; Nrxn1^flox/flox^; Nrxn2^flox/flox^; Nrxn3^flox/flox^*) (right panel) mice. Scale bar, 2 mm. **e**, Amount of protein in the cerebellum and forebrain of wild-type, control, and CGC-specific *Nrxn* TKO mice. The total proteins in mice from each genotype was quantified. **f**, Expression of NRXN proteins in wild-type, control, and CGC-specific *Nrxn* TKO mouse brains. Ten micrograms of homogenates prepared from the cerebellum and forebrain of mice from each genotype were separated by SDS-PAGE, followed by western blotting with anti-pan NRXN and β-actin antibodies, respectively. **g**, Quantification of the NRXN signals in **f**. WT, wild-type mice; CTL, control mice; TKO, CGC-specific *Nrxn* TKO mice. All values represent the mean ± s.e.m., n = 3 each from three animals. *** *P* < 0.001, ** *P* < 0.01, * *P* < 0.05, one-way ANOVA (*F* (2, 8) = 56.1, *P* = 1.3 × 10^−4^ in **e**; *F* (2, 8) = 16.7, P = 3.5 × 10^−3^ in **g**), followed by *post hoc* Tukey’s test (WT vs. TKO, *P* = 2.0 × 10^−4^, CTL vs. TKO, *P* = 2.7 × 10^−4^ in **e**; WT vs. TKO, *P* = 0.015, CTL vs. TKO, *P* = 0.0034 in **g**).

The size of the cerebellum of CGC-specific *Nrxn* TKO mice was smaller than that of control mice (Fig. 1d), and their total amount of cerebellar protein was reduced to approximately 50% of that of wildtype or control floxed *Nrxn (fNrxn*) mice (Fig. 1e). There was no substantial difference in that of forebrain proteins among genotypes. To examine the deletion of NRXN proteins in CGC-specific *Nrxn* TKO mice, we performed a western blot analysis using a pan anti-NRXN antibody. Protein bands of approximately 160–200 kDa, corresponding to the size of α-NRXNs^18^, were detected in cerebellar homogenates prepared from control *fNrxn* and wild-type mice. Conversely, the 90–100 kDa bands corresponding to the size of β-NRXNs^18^ were not clearly detected in these mice. In CGC-specific *Nrxn* TKO mice, these signals were reduced to 38% of those detected in control mice (Fig. 1f,g). There was no substantial difference in the NRXN signals in forebrain homogenates among genotypes. We estimated an approximately 80% reduction of cerebellar NRXN proteins in CGC-specific *Nrxn* TKO mice (Fig. 1e,g).

### Loss of CGCs in CGC-specific *Nrxn* TKO mice

CGC-specific *Nrxn* TKO mice showed severe ataxic gait and could not walk along a straight line using regular steps as control mice did. These mutant mice showed poor performance and no improvement in the constant-speed rotarod test (Supplementary Figure 2). To examine the formation of the cerebellar circuit, we performed an immunohistochemical analysis using antibodies against the vesicular glutamate transporter 1 (VGluT1), VGluT2, and calbindin. In the cerebellum, VGluT1 and VGluT2 are predominantly expressed in PF and CF terminals, respectively^19^. The parasagittal cerebellar sections prepared from 8-week-old CGC-specific *Nrxn* TKO mice were much smaller than those from control mice (Fig. 2a). The punctate staining signals for VGluT2 in the molecular layer were significantly increased in CGC-specific *Nrxn* TKO mice compared with control mice. In contrast, the staining signals for VGluT1 were decreased in the mutant mice compared with control mice (Fig. 2b,c). The VGluT1 signals in the mutant mice were different in each lobule, with the most prominent reduction detected in lobule IX (Fig. 2a,b). Unexpectedly, the DAPI staining signals were dramatically decreased in the GC layer of CGC-specific *Nrxn* TKO mice. The layered GC structure seemed to have disappeared in lobule IX of the mutant mice (Fig. 2b). Prominent reduction of VGluT1 signals in the molecular layer and of DAPI signals in the GC layer suggested the loss of CGCs in the mutant mice.

**Fig. 2.**
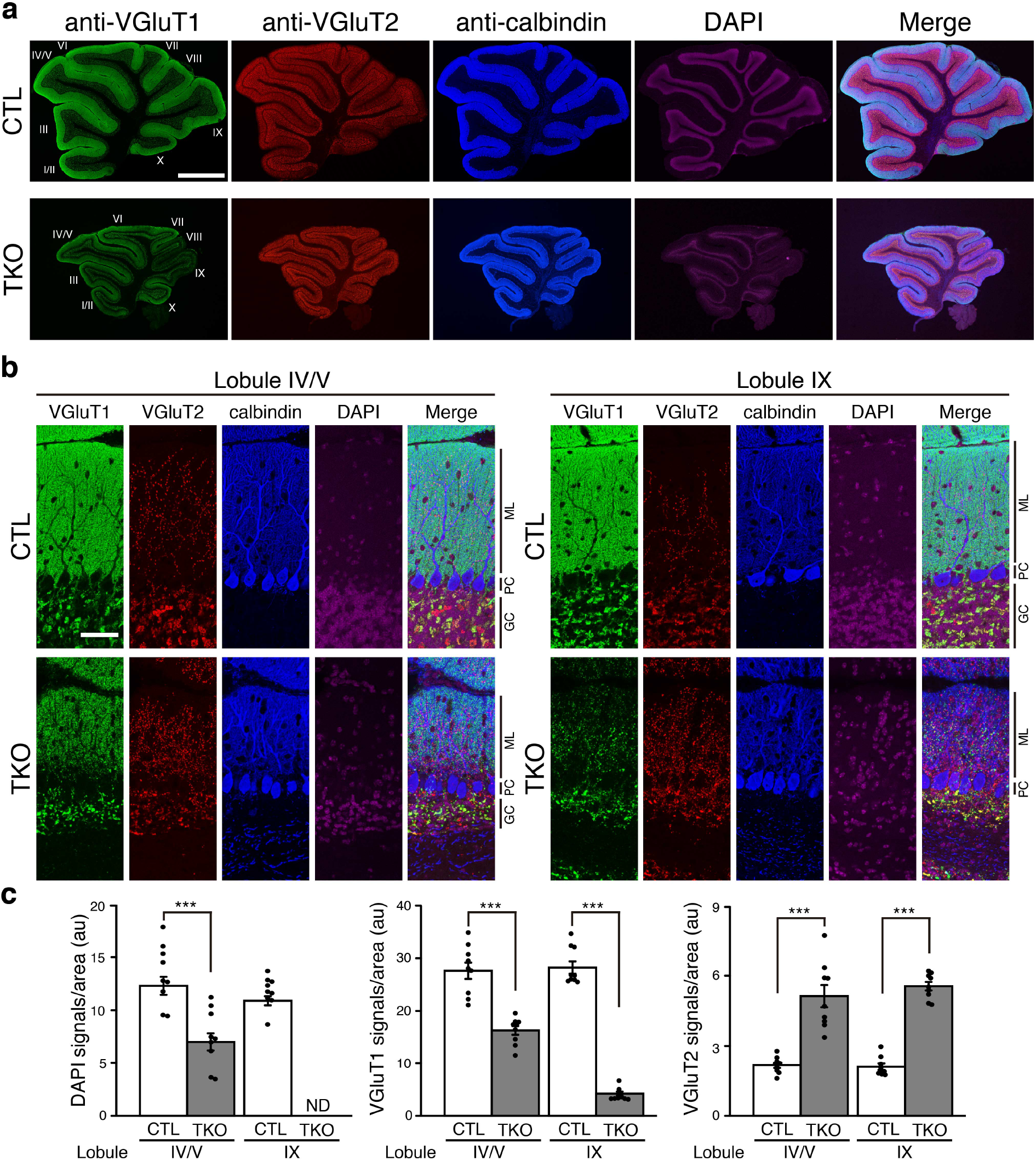
Immunohistochemical analysis of the cerebella of control and CGC-specific *Nrxn* TKO mice. **a**, The parasagittal cerebellar section from 8-week-old control and CGC-specific *Nrxn* TKO mice were stained with antibodies against VGluT1 (green), VGluT2 (red), calbindin (blue), and DAPI (purple), respectively. The images were merged (right panel). **b**, Enlarged images of the cerebellar cortex in **a**. The cerebellar lobules IV/V and IX are shown. ML, molecular layer; PC, Purkinje cell layer; GC, granule cell layer. **c**, Quantification of DAPI, VGluT1, and VGluT2 signals in **b**. The DAPI signal in the GC layer and the VGluT1 and VGluT2 signals in the molecular layer were quantified, respectively. All values represent the mean ± s.e.m., n = 6 each from three animals. *\*\*\*P* < 0.001, two-tailed Student’s *t*-test (DAPI, CLT vs. TKO in lobules IV/V, *P* = 3.6 × 10^−4^; CLT vs. TKO in lobule IX, *P* = 4.1 × 10^−7^; VGluT1, CLT vs. TKO in lobules IV/V, *P* = 3.0 × 10^−5^; CLT vs. TKO in lobule IX, *P* = 1.4 × 10^−12^; VGluT2, CLT vs. TKO in lobule IX, *P* = 1.7 × 10^−5^; and CLT vs. TKO in lobule IX, *P* = 1.8 × 10^−10^). CTL, control mice; TKO, CGC-specific *Nrxn* TKO mice; ND, not determined. Scale bars, 1 mm in **a** and 50 μm in **b**.

### NRXNs are essential for the survival of CGCs

In the mouse, the precursors of CGCs proliferate at the surface of the developing cerebellum and migrate inwardly to form the internal granule layer. This process ends by postnatal day 20 (P20), when cerebellar circuit formation is completed^20^. To examine whether the loss of CGCs in CGC-specific *Nrxn* TKO mice was caused by developmental defects, we analyzed the changes of the cerebellar structures with age. In the CGC-specific *Nrxn* TKO mice, the size of the cerebellum decreased gradually from 3 to 11 weeks of age (Fig. 3a). In lobules IV/V of 3-week-old mutant mice, the density of CGCs was comparable to that observed in control mice, but gradually decreased in as the animals aged. The well-defined layered structure of CGCs had almost completely disappeared at 11 weeks of age (Fig. 3b,c). Moreover, in lobule IX, the density of CGCs was decreased to 22% of that of control mice at 3 weeks of age, and the well-defined layered structure of CGCs had almost completely disappeared at 8 weeks of age (Fig. 3b,c). Conversely, the densities of PCs were increased in the mutant mice (Fig. 3b,d). An electron microscopic analysis revealed that the densities of PF-PC synapses were unchanged and decreased in lobules IV/V and IX of 4-week-old mutant mice, respectively. Consistent with the time course of CGC densities, the densities of PF-PC synapses were decreased from 4 weeks to 8 weeks in both lobules of the mutant mice (Supplementary Figure 3). These results suggest that CGCs were decreased in young mutant mice, whereas surviving CGCs formed synapses with PCs and died at a later stage. The differences in the time course of the density of CGCs among lobules were well matched with the developmental time course of Cre activity in *GluN2C^+/iCre^* mice, in which Cre-mediated recombination was first observed in lobules VIII and IX at P7 and gradually expanded to other lobules as the cerebellum developed (Supplementary Figure 4). Thus, the differences in the time course of the decrease in the density of CGCs is probably attributed to the difference in the time course of *Nrxn* deletion in each lobule. To examine whether the loss of CGCs in the CGC-specific *Nrxn* TKO mice is caused by apoptotic cell death, we performed TUNEL staining. The number of TUNEL-positive cells in the CGC layer was significantly increased in 3-week-old CGC-specific *Nrxn* TKO mice compared with control mice (Fig. 3e,f), suggesting the apoptotic death of CGCs. Collectively, these results indicate that NRXNs are essential for the survival of CGCs.

**Fig. 3.**
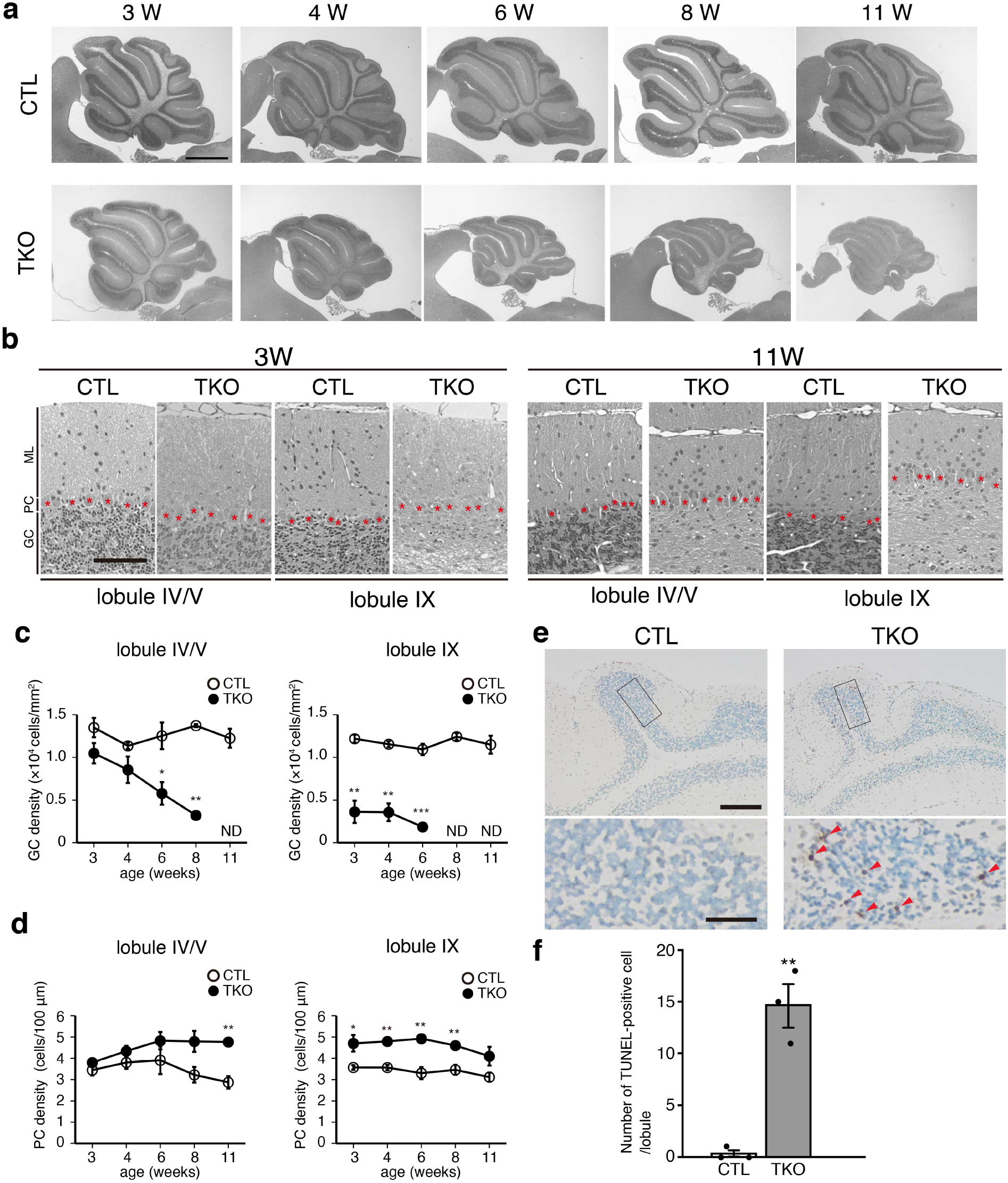
Progressive loss of CGCs in CGC-specific *Nrxn* TKO mice. **a**, Developmental neuroanatomical changes of the cerebella of control and CGC-specific *Nrxn* TKO mice. Parasagittal cerebellar sections prepared from 3, 4, 6, 8, and 11-week-old control and CGC-specific *Nrxn* TKO mice were stained with hematoxylin, respectively. **b**, Enlarged images of the cerebellar cortex of 3- and 11-week-old mice in **a**. The cerebellar lobules IV/V and IX are shown. The asterisks indicate the cerebellar PCs. ML, molecular layer; PC, Purkinje cell layer; GC, granule cell layer. **c**, Quantification of the developmental changes in CGC densities in control and CGC-specific *Nrxn* TKO mice. ND, not determined. All values represent the mean ± s.e.m., n = 9 each from three animals. \*\**P* < 0.01, \**P* < 0.05, two-way ANOVA (lobules IV/V, age × genotype interaction, *F* (3, 23) = 5.84, *P* = 0.0057; lobule IX, genotype effect, *F* (1, 17) = 180.7, *P* = 1.4 × 10^−8^), followed by *post hoc* twotailed Student’s *t*-test (CTL vs. TKO in lobules IV/V at 6 weeks, *P* = 0.021; CTL vs. TKO in lobules IV/V at 8 weeks, *P* = 2.6 × 10^−6^; CTL vs. TKO in lobule IX at 3, 4, and 6 weeks, *P* = 0.0035, 0.0019, and 2.1 × 10^−4^, respectively). **d**, Quantification of the developmental changes of cerebellar PC densities in control and CGC-specific *Nrxn* TKO mice. All values represent the mean ± s.e.m., n = 9 each from three animals. \*\**P* < 0.01, \**P* < 0.05, two-way ANOVA (lobules IV/V, genotype effect, *F* (1, 29) = 30.5, *P* = 2.1 × 10^−5^; lobule IX, genotype effect, *F* (1, 29) = 60.1, *P* = 1.9 × 10^−7^), followed by *post hoc* two-tailed Student’s *t*-test (control vs. TKO in lobules IV/V at 11 weeks, *P* = 0.005; control vs. TKO in lobule IX at 3, 4, 6, and 8 weeks, *P* = 0.048, 0.0064, 0.0069, and 0.011, respectively). **e**, TUNEL analysis of cerebella from control and CGC-specific *Nrxn* TKO mice at postnatal 3 weeks. The red arrowheads indicate the TUNEL-positive cells in the CGC layers. The enclosed images are shown at high resolution (lower panels). **f**, Quantification of TUNEL-positive cells in control and CGC-specific *Nrxn* TKO mice. All values represent the mean ± s.e.m., n = 3 from three animals each. \*\**P* < 0.01, two-tailed Student’s *t*-test (*P* = 0.0022). CTL, control mice; TKO, CGC-specific *Nrxn* TKO mice. Scale bars, 1 mm in **a**, 100 μm in **b**, 200 μm in **e** (upper panel), and 50 μm in **e** (lower panel).

Next, we examined whether NRXNs have functional redundancy regarding CGC survival. We generated CGC-specific *Nrxn1/2* double-knockout (DKO), *Nrxn2/3* DKO, and *Nrxn1/3* DKO mice (Supplementary Figure 5a). In contrast with CGC-specific *Nrxn* TKO mice, these mutant mice grew normally with no obvious ataxic gait, and showed apparently normal cerebellar structures (Supplementary Figure 5b). The DAPI signals in the GC layer and staining signals for VGluT1, VGluT2, and calbindin were indistinguishable from those detected in control mice (Supplementary Figure 5c). These results suggest that NRXN1, NRXN2, and NRXN3 are all expressed in CGCs and have functional redundancy, at least regarding CGCs survival. Finally, deletion of all NRNXs results in CGC death.

### PF–PC synapse-independent regulation of CGC survival by NRXNs

Next, we investigated whether the loss of CGCs in CGC-specific *Nrxn* TKO mice was caused by dysfunction and/or loss of PF-PC synapses. For this, we used cultured CGCs, as they form no synapses among them and the majority of presynaptic-like structures with presynaptic vesicular profiles are not opposed to definite postsynaptic structures^21–23^, whereas functional synapses may be formed only in a small portion of CGCs, as synaptic transmission has been observed in CGCs^24^. The cultured CGCs that were prepared from *fNrxn* pups (termed cultured *fNrxn* CGCs) were used to analyze the effects of *Nrxn* TKO on CGC survival. In our cultured condition, approximately 96% of cells were NeuN-positive CGCs (Supplementary Figure 6a). For conditional *Nrxn* TKO, infection with lentivirus-iCre was used (Supplementary Figure 6b). Six days after infection, the numerous punctate signals for NRXNs along the axons of CGCs had disappeared almost completely (Fig. 4a), suggesting the successful knockout of all NRXNs. Infection with lentivirus-iCre in the cultured *fNrxn* CGCs resulted in a 13.3%, 59.7%, and 87.3% reduction in the number of CGCs compared with that detected in control cultures at days in vitro (DIV) 4, 7, and 10, respectively (Fig. 4b,c). Co-infection with a lentivirus expressing NRXN1β(+S4) or NRXN1β(–S4) restored the defects of CGC viability. These results suggest that NRXNs regulate the viability of cultured CGCs and that expression of NRXN1β, one of the NRXN variants, is sufficient for CGC survival. Moreover, based on the characteristic features of cultured CGCs, these results also suggest that NRXNs regulate the survival of cultured CGCs independent of the contact with postsynaptic structures.

**Fig. 4.**
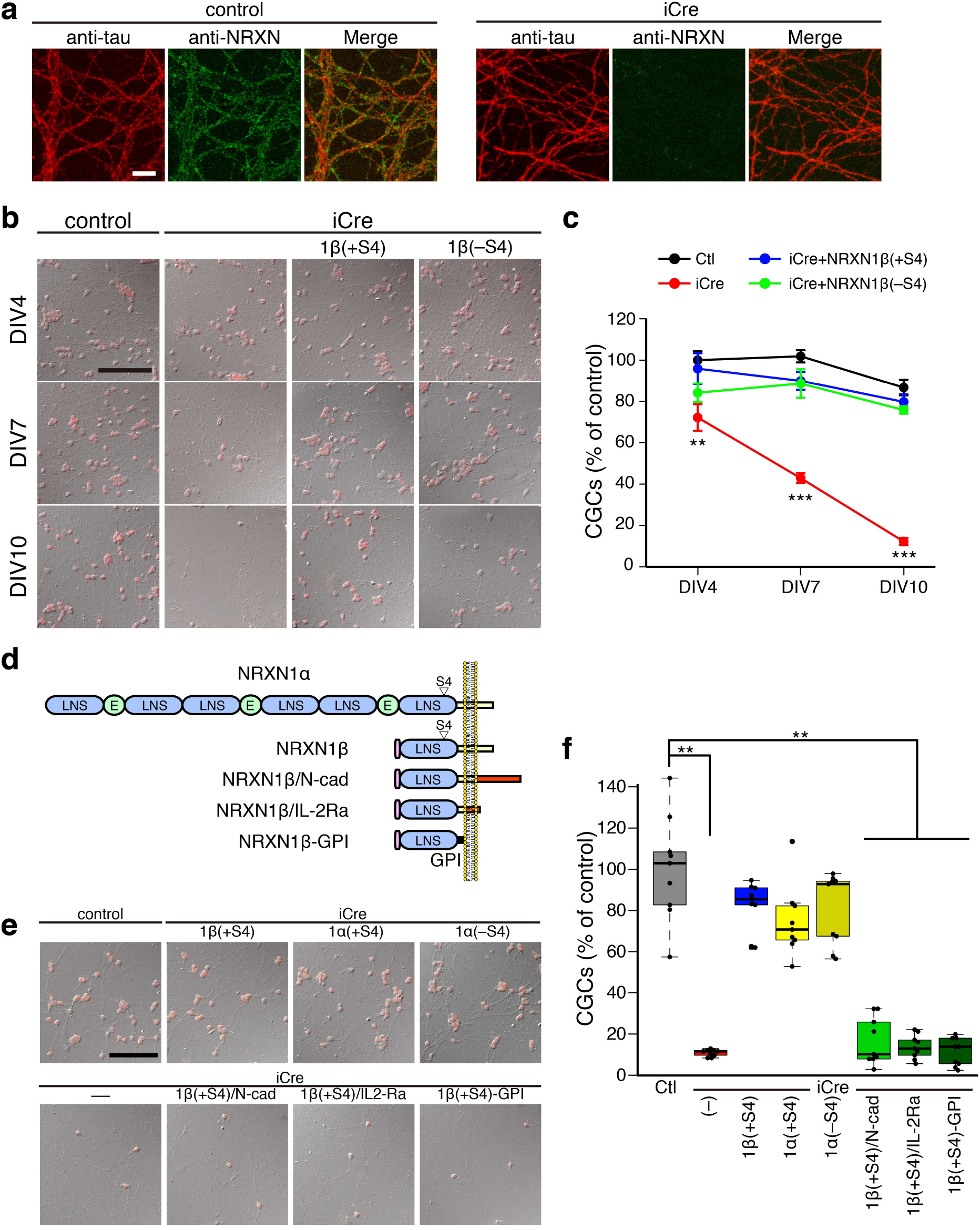
NRXNs are essential for cultured CGC survival. **a**, Lentivirus-iCre mediated *Nrxn* TKO in cultured CGCs. The cultured *fNrxn* CGCs were infected with lentivirus-iCre at DIV1 and stained with antibodies against tau and NRXNs at DIV7. **b**, Effects of *Nrxn* TKO on cultured CGC survival. The cultured *fNrxn* CGCs were infected with lentivirus-iCre alone or together with HA-NRXN1β(+S4) or HA-NRXN1β(−S4). The cells were stained with an antibody against NeuN (red), and differential interference contrast (DIC) merged images are shown. **c**, Quantification of the number of NeuN-positive CGCs in **b** (n = 6 each). \*\*\**P* < 0.001, \*\**P* < 0.01; two-way ANOVA (time × group interaction, *F* (6, 71) = 7.06, *P* = 1.0 × 10^−5^), followed by *post hoc* Tukey’s test (CTL vs. iCre at DIV4, 7, and 10 *P* = 0.015, 6.8 × 10^−8^, and 1.4 × 10^−12^, respectively). **d**, Schematic structures of the NRXN1α, NRXN1β, and NRXN1β mutants. **e**, Effects of the expression of NRXN1α and NRXN1β and its NRXN1β mutants on *Nrxn* TKO CGC survival. The cultured *fNrxn* CGCs were infected with iCre alone or together with *Nrxn* variants or mutants. The cells were stained with an antibody against NeuN (red) at DIV 10, and DIC merged images are shown. **f**, Quantification of the number of NeuN-positive CGCs in **e** (n = 9 each). The horizontal line in each box indicates the median, the box shows the interquartile range (IQR), and the whiskers are the 1.5× IQR. \*\**P* < 0.01; Steel-Dwass test (Ctl vs. iCre, NRXN1β(+S4)/N-cad, NRXN1β(+S4)/IL2-Ra, and NRXN1β(+S4)-GPI, *P* = 0.0082, 0.0083, 0.0084, and 0.0083, respectively). Scale bars, 10 μm in **a** and 100 μm in **b** and **e**.

### The C-terminal region of NRXN1β is essential for the survival of cultured CGCs

NRXNs have a long α-NRXN and a short β-NRXN variant produced by two alternative promoters (Fig. 4d). Thus, we examined whether α-NRXNs have the same function regarding the survival of cultured CGCs. Lentivirus-iCre mediated knockout of all NRXNs resulted in a decrease in the number of CGCs, which was restored by co-infection with NRXN1β(+S4), NRXN1α(+S4), or NRXN1α(−S4) (Fig. 4e,f). There were no substantial differences in cell survival among NRXN1β(+S4), NRXN1α(+S4), and NRXN1α(−S4) co-infected cultures, suggesting that NRXN1β and NRXN1α have the same function, at least regarding cultured CGC survival. The extracellular regions are not identical among the NRXN1 variants, but share some sequences. Conversely, the transmembrane and cytoplasmic regions are identical among them. Thus, we speculated that the transmembrane and/or cytoplasmic region of NRXN1 are responsible for cultured CGC survival. To examine this hypothesis, we constructed three NRXN1β(+S4) mutants: (1) the cytoplasmic region of NRXN1β(+S4) was replaced with that of N-cadherin (NRXN1β(+S4)/N-cad), (2) the extracellular domain of NRXN1β(+S4) was fused to the transmembrane and cytoplasmic regions of interleukin-2 receptor-alpha (IL-2Ra) (NXN1β(+S4)/IL-2Ra); and (3) the extracellular region of NRXN1β(+S4) was fused to GPI (NRXN1β(+S4)-GPI) (Fig. 4d). The reduction in the number of CGCs in iCre-infected cultures was not rescued by these mutants (Fig. 4e,f). These results suggest that NRXNs regulate the viability of cultured CGCs through their C-terminal region.

### Non-cell-autonomous factors are sufficient for the survival of cultured *Nrxn* TKO CGCs

To confirm further the role of NRXNs in cultured CGC survival, the cultured *fNrxn* CGCs were sparsely transfected with the expression vector for EGFP with or without that for iCre. In control EGFP-transfected *fNrxn* CGCs, numerous punctate staining signals for NRXNs were detected along the axons (Supplementary Figure 7a). These signals could not be detected in EGFP and iCre-transfected *fNrxn* CGCs, suggesting knockout of all NRXNs. Unexpectedly, the number of EGFP-positive CGCs was comparable between control and iCre-transfected *fNrxn* CGCs (Supplementary Figure 7b,c). This result was entirely different from the results obtained for the lentiviral-mediated *Nrxn* TKO (Fig. 4b-f). As one of the major differences between these two experiments was the population of *Nrxn* TKO cells used in the cultures (Supplementary Figures 6 and 7), we hypothesized that a non-cell-autonomous factor affects CGC survival. To examine whether cultured CGC survival is regulated non-cell-autonomously, we prepared a mixed culture of wild-type and *Nrxn* TKO CGCs (Fig. 5a). The co-culture of EGFP-labeled *Nrxn* TKO CGCs with non-EGFP-labeled *Nrxn* TKO CGCs led to a significant decrease in the numbers of both EGFP labeled and non-labeled cells compared with those in control cultures. Co-culture with non-EGFP-labeled wild-type CGCs restored the number of EGFP-labeled *Nrxn* TKO CGCs to the same level as that detected in the control cultures (Fig. 5b,c). These results suggest that non-cell-autonomous factors and/or factors from wild-type CGCs are sufficient for the survival of *Nrxn* TKO CGCs.

**Fig. 5.**
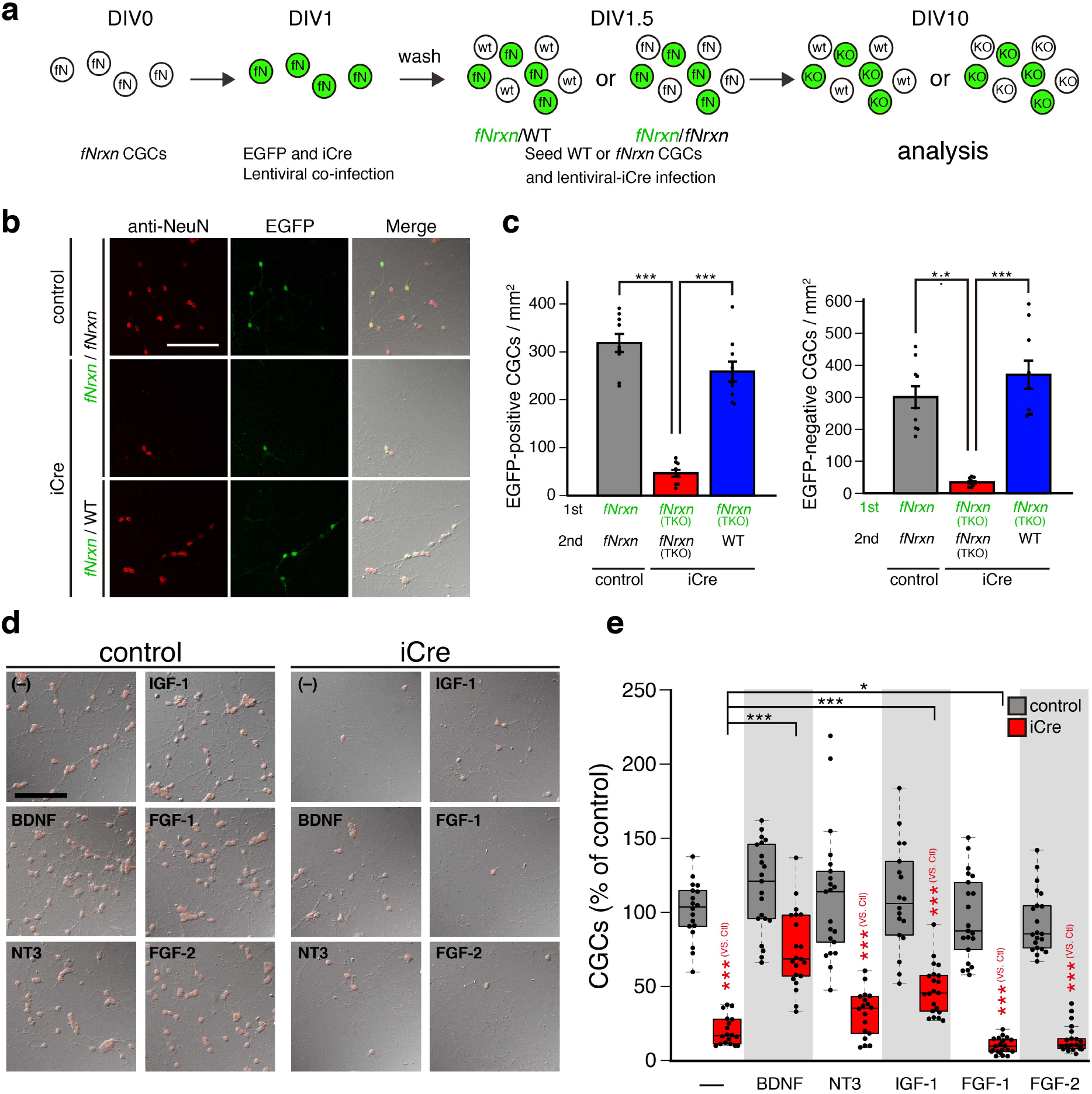
Non-cell-autonomous factors rescue the cell survival defects in cultured *Nrxn* TKO CGCs. **a**, Schema used for the analysis of the effects of the context on the survival of *Nrxn* TKO CGCs. In this assay, EGFP-labeled *Nrxn* TKO CGCs were cultured in an ambient environment of wild-type or *Nrxn* TKO CGCs. Cultured *fNrxn* CGCs were infected with a lentivirus expressing iCre and EGFP at DIV1. After extensive washing, *fNrxn* CGCs or wild-type CGCs were seeded at DIV1.5, followed by infection of these cells with lentivirus-iCre. fN, *fNrxn* CGCs; wt, wild-type CGCs; KO, *Nrxn* TKO CGCs. **b**, Effects of the context on the survival of *Nrxn* TKO CGCs. The cells were stained with an antibody against NeuN (red) at DIV10, and DIC merged images are shown. **c**, Number of EGFP-negative or EGFP-positive CGCs. All values represent the mean ± s.e.m., n = 9 each. \*\*\**P* < 0.001, one-way ANOVA (EGFP-positive CGCs, *F* (2, 26) = 64.7, *P* = 2.2 × 10^−10^; EGFP-negative CGCs, *F* (2, 26) = 27.1, *P* = 7.0 × 10^−7^), followed by *post hoc* Tukey’s test (EGFP-positive CGCs, fNrxn/fNrsn-iCre vs. fNrxn/fNrsn-contrl and fNrxn/WT-iCre, *P* = 5.4 × 10^−9^ and 4.0 × 10^−8^, respectively; EGFP-negative CGCs, fNrxn/fNrsn-iCre vs. fNrxn/fNrsn-contrl and fNrxn/WT-iCre, P = 3.2 × 10^−5^ and 9.7 × 10^−7^, respectively). **d**, Effects of exogenous neurotrophic factors on the survival of *Nrxn* TKO CGCs. The indicated neurotrophic factors were applied in control or *Nrxn* TKO CGCs. The cells were stained with antibody against NeuN (red), and DIC merged images are shown. **e**, Number of NeuN-positive CGCs in control and *Nrxn* TKO CGC with or without the indicated neurotrophic factors (n = 18-21). The horizontal line in each box indicates the median, the box shows the interquartile range (IQR), and the whiskers are the 1.5× IQR. \*\*\**P* < 0.001, n = 18 (control, iCre, and iCre+NT3) and 21 (others), Steel-Dwass test (control vs. iCre, iCre+NT3, iCre+IGF-1, iCre+FGF-1, and iCre+FGF-2, *P* = 1.9 × 10^−5^, 2.3 × 10^−5^, 2.6 × 10^−5^, 6.5 × 10^−6^, and 6.6 × 10^−6^, respectively; iCre vs. iCre+BDNF, iCre+IGF-1, and iCre+FGF-1, *P* = 1.4 × 10^−5^, 2.0 × 10^−4^, and 0.021, respectively). Scale bars, 100 μm.

Given that cultured CGCs do not form synapses among them, the paracrine action of neurotrophic factors would be one of the candidates for supporting *Nrxn* TKO CGC survival non-cell-autonomously in the mixed culture. Thus, to examine the effects of neurotrophic factors on the survival of cultured *Nrxn* TKO CGCs, we applied BDNF, neurotrophin (NT3), insulin-like growth factor-1 (IGF-1), fibroblast growth factor-1 (FGF-1), or FGF-2 extracellularly. The application of BDNF and IGF-1 significantly increased the cell viability of *Nrxn* TKO CGCs (Fig. 5d,e). In contrast, NT3, FGF-1, and FGF-2 had no positive effect on the survival of these cells. Among the neurotrophic factors tested here, BDNF was the most effective regarding the survival of cultured *Nrxn* TKO CGCs. These results raise the possibility that neurotrophic factor secretion is impaired in these cells.

### Cultured *Nrxn* TKO CGCs have an impaired neurotrophic factor secretory machinery in axons

To examine whether the neurotrophic factor secretory machinery is impaired in *Nrxn* TKO CGCs, we used a super-ecliptic pHluorin-tagged BDNF (BDNF-pHluorin). The *fNrxn* CGCs were sparsely transfected with BDNF-pHluorin with or without iCre, and the BDNF-pHluorin secretion was quantified by monitoring its fluorescence increase upon exocytic secretion to the extracellular space. In control cultures, the number of BDNF-pHluorin puncta along the axons was significantly increased after the application of 50 mM KCl, suggesting the BDNF-pHluorin secretion from axons. In contrast, that in *Nrxn* TKO CGCs was not increased by the application of KCl at high concentration (Fig. 6a,b). Expression of NRXN1β(+S4) restored the high-KCl-induced BDNF-pHluorin fluorescence increase. Conversely, the expression of the chimeric mutant NRXN1β(+S4)/N-cad failed to rescue the fluorescence increase (Fig. 6a,b). These results suggest that the machinery of axonal neurotrophic factor secretion is impaired in *Nrxn* TKO CGCs and that it requires the C-terminal region of NRXNs.

**Fig. 6.**
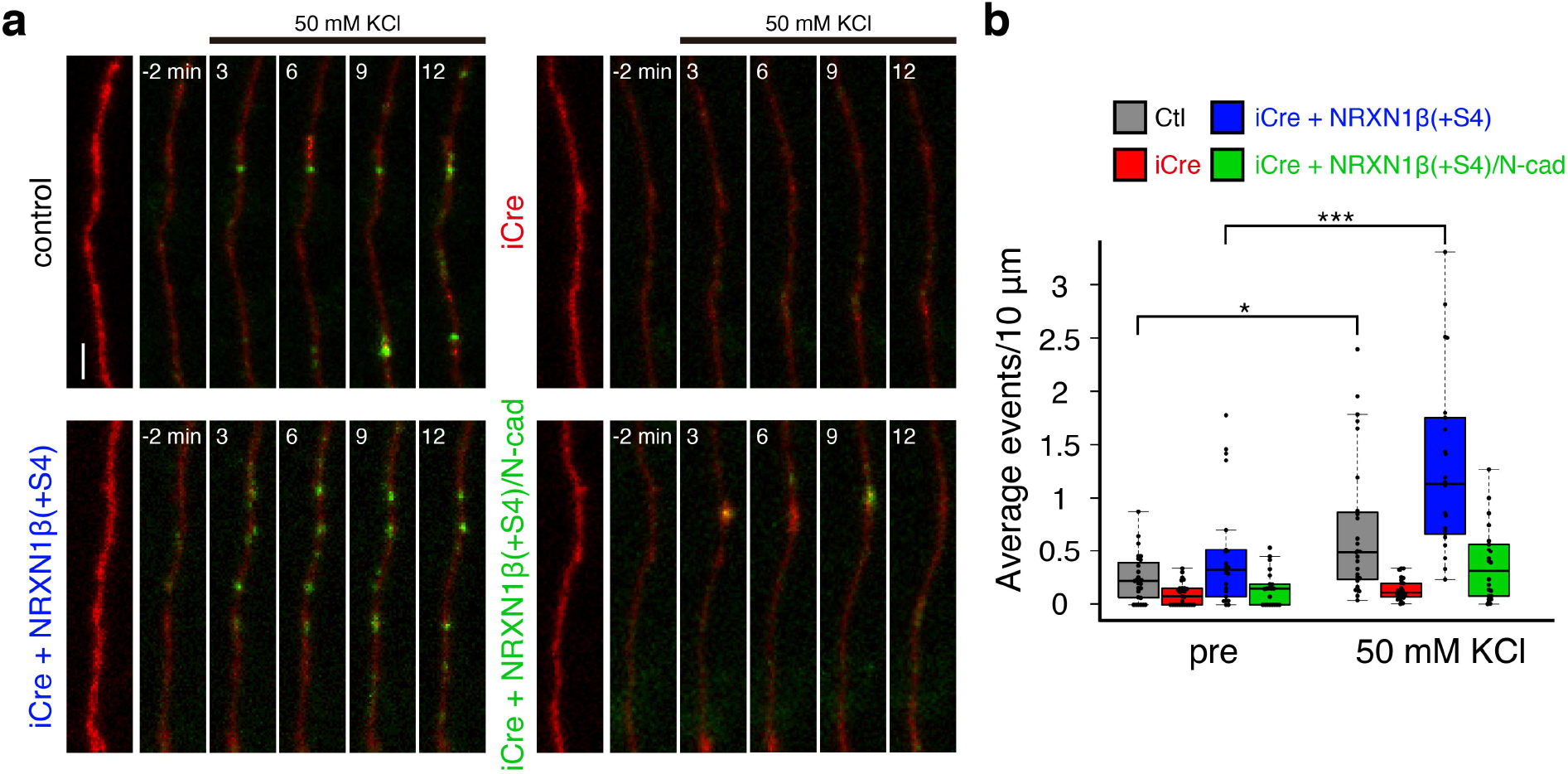
Impairment of activity-induced axonal BDNF secretion in cultured *Nrxn* TKO CGCs. **a**, Time-lapse images of BDNF-pHluorin induced by stimulation with 50 mM KCl. The cultured *fNrxn* CGCs were transfected with TagRFP together with the indicated expression vectors. Images were acquired 2 min before stimulation and every 3 min after stimulation. The bright images of TagRFP are shown on the left. Scale bar, 2 μm. **b**, BDNF-pHluorin fluorescence events before and after stimulations. Average events before and after stimulation were quantified in control, iCre, iCre and NRXN1β(+S4), and iCre and NRXN1β(+S4)/N-cad, respectively. The horizontal line in each box indicates the median, the box shows the interquartile range (IQR), and the whiskers are the 1.5× IQR. \*\*\**P* < 0.001, *\*P* < 0.05, n = 27 (CTL and iCre) and n = 22 (iCre+ NRXN1β(+S4) and iCre+ NRXN1β(+S4)/N-cad), Steel-Dwass test (pre Ctl vs. after Ctl, *P* = 0.047; pre iCre vs. after iCre + NRXN1β(+S4), *P* = 8.8 × 10^−8^).

### Cultured *Nrxn* TKO CGCs exhibit impaired action potential-induced Ca^2+^ influx into presynaptic-like structures

Neurotrophic factors are stored in dense-core vesicles, and exocytotic vesicular secretion is triggered by membrane depolarization-induced Ca^2+^ influx^25^. In cultured CGCs, activity-dependent Ca^2+^ influx and exocytosis occur at the site of axonal presynaptic-like structures with synaptic vesicle-like assemblies^21,22^. Thus, to examine the membrane-depolarization-induced Ca^2+^ influx in the presynaptic-like structure of CGCs, we constructed vesicle-associated membrane protein-2 (VAMP2) fused with the fluorescent calcium sensor GCaMP6s^26^ at its N terminus (GCaMP6s-VAMP2). A previous study showed that GCaMP5G-synaptobrevin-2/VAMP2 is targeted to the presynaptic terminal and can be used for the analysis of presynaptic Ca^2+^ transients^4^. The cultured CGCs were sparsely transfected with GCaMP6s-VAMP2 and TagRFP, and stained with antibodies against the active zone protein Bassoon and GCaMP6s. The majority of GCaMP6s-VAMP2 signals were well merged with Bassoon signals (Fig.7a). Electrical-field stimulation evoked robust Ca^2+^-induced fluorescence transients in the axons of GCaMP6s-VAMP2-transfected CGCs (Fig. 7b-d). To examine Ca^2+^ transients in the axon of *Nrxn* TKO CGCs, cultured *fNrxn* CGCs were transfected with GCaMP6s-VAMP2 with or without iCre. Fluorescence transients were elicited by stimulation, and relative changes in fluorescence intensity were quantified. In control cultures, fluorescence intensities in the axons were elicited by stimulation at 50 Hz (2–80 stimuli), which were saturated after 25 stimuli (Fig. 7e,f). Fluorescence transients were blocked by tetrodotoxin, suggesting that the Ca^2+^ influx was induced by action potential stimulation. In *Nrxn* TKO CGCs, the action potential-induced fluorescence transients were decreased, suggesting that action potential-induced Ca^2+^ influx into presynaptic like structure is impaired. Decreased fluorescence transients were restored by co-transfection with NRXN1β(+S4), but not the chimeric mutant NRXN1β(+S4)/N-cad (Fig. 7e,f).

**Fig.7.**
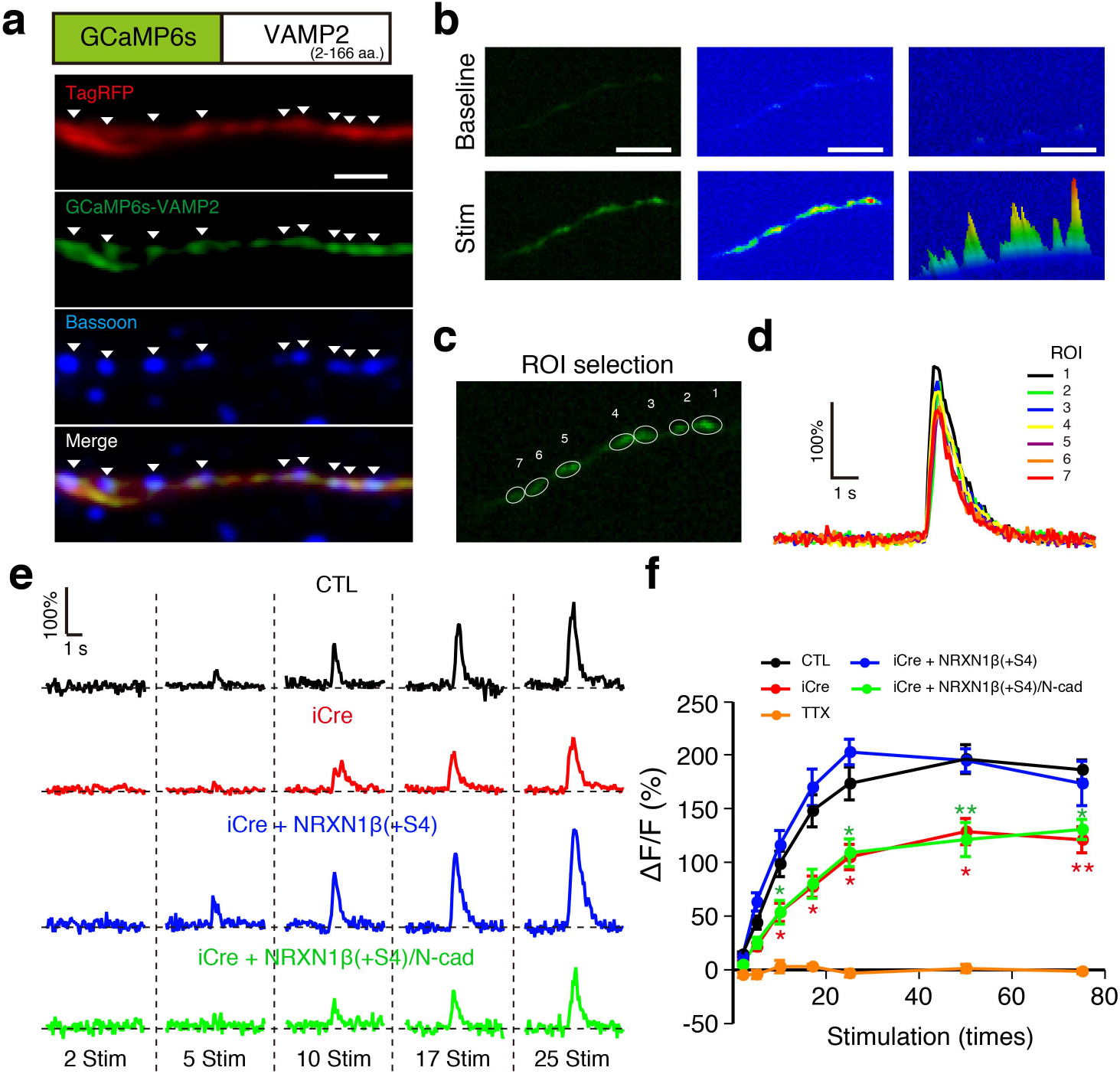
Impairment of the action potential-induced Ca2^+^ influx into axonal varicosities of cultured *Nrxn* TKO CGCs. **a**, Localization of GCaMP6s-VAMP2 in cultured CGCs. Schematic representation of the GCaMP6s-VAMP2 fusion protein (top). The cultured CGCs were transfected with TagRFP and GCaMP6s-VAMP2, and stained with an antibody against Bassoon and EGFP. The arrowheads indicate GCaMP6s-VAMP2 signals merged with Bassoon signals. **b**, **c**, Representative Ca^2+^ imaging experiment. The cultured CGCs were transfected with TagRFP and GCaMP6s-VAMP2 at DIV3. The imaging experiments were performed at DIV7 or DIV8. Representative images of the fluorescence responses before and after 75 stimuli applied at 20 Hz are shown in **b**. Left panels, fluorescence images; middle panels, false-colored images; right panels, 3D surface plot of the false-colored images in **b**. The regions of interest (ROIs) used in the quantitative analysis are shown in **c**. The fluorescence traces of GCaMP6s-VAMP2 fluorescence in **c** are shown in **d**. **e**, Effect of *Nrxn-TKO* CGC on action-potential-induced Ca^2+^ influx in the axons. The cultured *fNrxn* CGCs were transfected with iCre alone or together with NRXN1β(+S4) or C-terminal chimeric mutant NRXN1β(+S4)/N-cad. Representative fluorescence traces to 2, 5, 10, 17, and 25 stimuli applied at 20 Hz are shown. **f**, Summary of the results (n = 10-18 neurons) presented in **e**. All values represent the mean ± s.e.m. \*\**P* < 0.01, \**P* < 0.05, n = 20 (CTL), n = 13 (iCre), n = 12 (iCe+NRN1β(+S4), and n = 13 neurons (iCe+NRN1β(+S4)/N-cad), two-way repeated ANOVA (time × group interaction, *F* (18, 405) = 5.01, *P* = 4.6 × 10^−10^), followed by *post hoc* Tukey’s test (CTL vs. iCre at 2, 5, 10, 17, 25, 50, and 70 min, *P* = 0.621, 0.055, 0.04, 0.004, 0.005, 0.004, and 0.002, respectively; CTL vs. iCe+NRN1β(+S4)/N-cad at 2, 5, 10, 17, 25, 50, and 70 min, *P* = 0.035, 0.151, 0.041, 0.006, 0.009, 0.001, and 0.01, respectively). Scale bars, 1 μm in **a** and 5 μm in **b**.

### NRXNs are essential for the formation of presynaptic-like structures in cultured CGCs axons

The presynaptic-like structures in the axons of CGCs have vesicular profiles that are similar to those of synaptic vesicle structures, in which presynaptic proteins, such as the active zone protein Bassoon, VGluT1, and synapsin, are accumulated^21–23^. Thus, we examined the formation of presynaptic-like structures in *Nrxn* TKO CGCs. *fNrxn* CGCs were sparsely transfected with EGFP with or without iCre, and immunostained with antibodies against Bassoon, synapsin, and VGluT1, respectively. *Nrxn* TKO CGCs showed a significant decrease in the densities of punctate staining signals for Bassoon, synapsin, and VGluT1 along the axons compared with those detected in control CGCs (Fig. 8a-f). Furthermore, the intensities of those signals were significantly decreased in *Nrxn* TKO CGCs (Fig. 8a-c and 8g-i). Co-expression of NRXN1β(+S4) restored both the densities and intensities of Bassoon, synapsin, and VGluT1 puncta. Conversely, the chimeric mutant NRXN1β(+S4)/N-cad failed to restore these defects (Fig. 8a-i). These results suggest that NRXNs regulate the formation of axonal presynaptic-like structures having presynaptic proteins through their C-terminal region, independent of PC contacts.

**Fig. 8.**
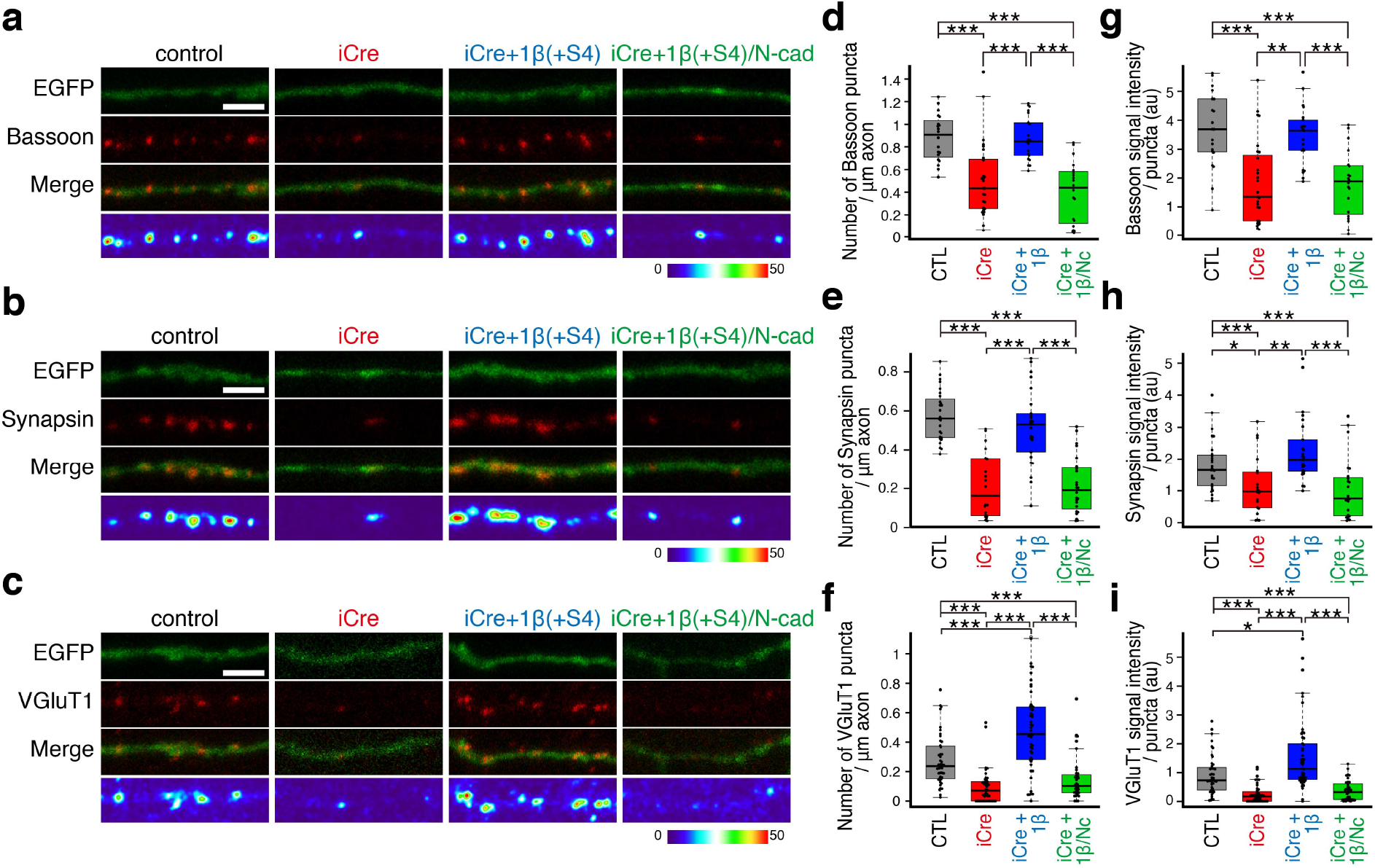
Cultured *Nrxn* TKO CGCs exhibit defects in the formation of presynaptic elements. **a-c**, Effect of *Nrxn* TKO on the formation of presynaptic elements. The cultured *fNrxns* were transfected with iCre and EGFP or together with NRXN1 β(+S4) or C-terminal chimeric mutant NRXN 1 β(+S4)/N-cad at DIV3 and immunostained with antibodies against Bassoon (**a**), synapsin (**b**), and VGluT1 (**c**) at DIV7, respectively. False-colored images of fluorescence signal intensities are shown in **a**-**c** (bottom panels). **d**-**f**, Densities of Bassoon (**d**), synapsin (**e**), and VGluT1 (**f**) puncta along the axons of CGCs in **a**-**c**. *\*\*\*P* < 0.001, Steel-Dwass test (CTL vs. iCre and iCre+1β(+S4)/Nca-d, *P* = 4.3 × 10^−4^ and 2.5 × 10^−5^, respectively. iCre+1β(+S4) vs. iCre and 1β(+S4)/N-cad, *P* = 2.4 × 10^−4^ and 1.2 × 10^−5^, respectively in **d**; CTL vs. iCre and iCre+1β(+S4)/N-cad, *P* = 9.1 × 10^−7^ and 1.1 × 10^−7^, respectively. iCre+1β(+S4) vs. iCre and 1β(+S4)/N-cad, *P* = 6.1 × 10^−5^ and 9.4 × 10^−6^, respectively in **e**; CTL vs. iCre, iCre+1β(+S4), and iCre+1β(+S4)/N-cad, *P* = 5.3 × 10^−8^, 8.3 × 10^−4^, and 3.9 × 10^−4^, respectively; iCre+1β(+S4) vs. iCre and 1β(+S4)/N-cad, *P* = 1.9 × 10^−10^ and 4.1 × 10^−7^, respectively in **f**. **g**-**i**, Intensities of punctate staining signals for Bassoon (**g**), synapsin (**h**), and VGluT1 (**i**) along the axons of CGCs in **a**-**c**. *\*\*\*P* < 0.001, *\*\*P* < 0.01, *\*P* < 0.05; Steel-Dwass test (CTL vs. iCre and iCre+1β(+S4) N-cad, *P* = 6.2 × 10^−4^ and 7.9 × 10^−4^, respectively. iCre+1β(+S4) vs. iCre and 1β(+S4)/N-cad, *P* = 1.1 × 10^−4^ and 4.3 × 10^−4^, respectively in **g**; CTL vs. iCre and iCre+1β(+S4)/N-cad, *P* = 0.028 and 0.0057, respectively. iCre+1β(+S4) vs. iCre and 1β(+S4)/N-cad, *P* = 0.0013 and 1.4 × 10^−4^, respectively in **h**; CTL vs. iCre, iCre+1β(+S4), and iCre+1β(+S4)/N-cad, *P* = 2.4 × 10^−7^, 1.0 × 10^−4^, and 3.0 × 10^−4^, respectively; iCre+1β(+S4) vs. iCre and 1β(+S4)/N-cad, *P* = 1.2 × 10^−11^ and 4.9 × 10^−9^, respectively in **i**. n = 20, 27, 21, and 20 (CTL, iCre, iCre+1β(+S4), and 1β(+S4)/N-cad, respectively) in **d** and **g**; n = 27, 20, 25, and 24 (CTL, iCre, iCre+ 1β(+S4), and 1β(+S4)/N-cad, respectively) in **e** and **h**; n = 44, 44, 39, and 36 (CTL, iCre, iCre+ 1β(+S4), and 1β(+S4)/N-cad, respectively) in **f** and **i**. Scale bars, 2 μm. The horizontal line in each box indicates the median, the box shows the interquartile range (IQR), and the whiskers are the 1.5× IQR in **d**-**i**.

## DISCUSSION

In the cerebellum, the GluD2-Cbln1-NRXN ternary complex regulates PF–PC synapse formation^12^. The analysis of both conventional and conditional KO mice provided evidence that GluD2 and Cbln1 play an essential role in the formation and maintenance of PF-PC synapses and in cerebellar LTD^12–16^. However, the physiological role of NRXNs in CGCs has not been fully clarified. Here, we generated CGC-specific *Nrxn* KO mice and found that NRXNs organize the axonal neurotrophic factor secretory machinery and are thus essential for CGCs survival. We also discovered that NRXNs regulate the formation of presynaptic-like structures independent of postsynaptic ligands.

CGC-specific *Nrxn* TKO mice showed severe loss of CGCs (Fig. 2). A developmental analysis of CGC-specific *Nrxn* TKO mice cerebella suggested that NRXNs play a crucial role in the survival of CGCs (Fig. 3). These phenotypes were entirely different from those of conventional and conditional *GluD2* or *Cbln1* KO mice^12–16^. Recent studies of four different types of synapses in conditional *Nrxn* TKO mice, i.e., CF-PC synapse, synapses formed by parvalbumin- or somatostatin-positive interneurons on pyramidal layer 5 neurons in the medial prefrontal cortex, and the calyx of Held synapse, reported defects in synapse-specific functions, ranging from a decrease in synaptic density to impairments of synaptic transmission and action-potential-induced Ca^2+^ influx^8,9^. Intriguingly, the effect of the deletion of all NRXNs in CGCs was strikingly different from those effects. During development, defects in synaptic connection or functions often induce apoptotic neuronal cell death in the brain^27^. In fact, a reduction of the CGC layer (14% reduction) was observed in GluD2-deficient mice, in which PF-PC synapse number was decreased to approximately half of that observed in wild-type mice^14^. Several mutant mice, such as PC degeneration mice, cysteine protease cathepsin B and C double KO, and Zürich I prion protein (PrP) KO mice expressing a truncated PrP, exhibited an almost complete loss of cerebellar PC and a decreased number of CGCs; nevertheless, CGCs were still present in the inner GC layer^28–30^. These phenotypes were entirely different from our observation of almost complete loss of CGCs in CGC-specific *Nrxn* TKO mice (Fig. 3). Thus, complete loss of CGCs in the mutant mice cannot be ascribed to PF-PC synapse loss or dysfunction. To address this, we used cultured CGCs and showed the requirement of NRXNs for the survival of cultured CGCs (Fig. 4). Importantly, the cultured CGCs do not form synapses among them, and the axonal presynaptic-like structures containing presynaptic proteins lack contact with postsynaptic structures^21–23^. Thus, our results suggest that NRXNs regulate CGC survival in a non-synaptic manner. Consistent with this notion, functional and morphological defects were observed in cultured *Nrxn* TKO CGC axons (Figures 6–8).

It has been reported that neurotrophic factors support neuronal survival^25^. In the cultured *Nrxn* TKO CGCs, the defect of CGC survival was rescued by co-culture with wild-type CGCs or the application of the neurotrophic factors BDNF or (partially) IGF-1 (Fig. 5). In cultured *Nrxn* TKO CGCs, activity-dependent axonal BDNF-pHluorin secretion was impaired (Fig. 6). These results suggest that neurotrophic factors act in a autocrine and/or paracrine manner for the survival of cultured CGCs, and that one of the causes of *Nrxn* TKO CGC death should be the impairment of neurotrophic factor secretion from the axons of CGCs. Consistent with this notion, it is suggested that the depolarization-induced Ca^2+^ influx that is critical for CGC survival occurs in the axonal varicosity compartment, rather than the dendritic compartment^22^. Unlike axonal varicosities in the cultured CGCs, axonal varicosities of PF that make synapses in contact with the PCs are covered with glial processes *in vivo*. Conversely, PFs form bundles with honeycomb structure without glial processes^11^. Thus, in addition to autocrine action at PF-PC synapses, autocrine and/or paracrine action at axons may work *in vivo* to support CGC survival. Autocrine regulation of neuronal survival by neurotrophic factors was also reported in cultured dorsal root ganglion and hippocampal neurons^31,32^. In our experimental condition, BDNF was most effective on the survival of *Nrxn* TKO CGCs (Fig. 4e,f). BDNF and its receptor, TrkB, are reportedly expressed in cultured CGCs^33^. Previous studies showed that BDNF, NT-3, FGF-2, and IGF-1 enhance the survival of cultured CGCs^33–36^, albeit with variable effects, presumably because of the developmental stage of the CGCs. In the rodent cerebellum, the BDNF and NT-3 mRNAs are expressed in CGCs, which also express TrkB and TrkC^37–39^. Several studies have suggested that BDNF and NT-3 are secreted from PF terminals and act in an anterograde or autocrine fashion^40–43^. Conversely, IGF-1 is predominantly expressed in cerebellar PCs, while its receptor is expressed in both cerebellar PCs and CGCs^44–46^. It is thought that IGF-1 secreted by PC is taken up by CGCs^34,44^. Therefore, it is possible that multiple factors both from PFs and PCs sustain CGC survival *in vivo*.

To date, several molecules that regulate neurotrophic factor release have been identified. BDNF, which is one of the most studied neurotrophic factors, is stored in dense-core vesicles and its release is normally triggered by membrane depolarization induced Ca^2+^ influx through voltage-gated Ca^2+^ channels^25^. However, there are several unsolved questions about how its release site is constructed and functioned. In the cultured CGCs, we showed that action-potential-induced Ca^2+^ influx occurred at the axonal presynaptic-like structure (Fig. 7). Consistently, previous studies showed that high-KCl-induced Ca^2+^ influx and exo-endocytosis non-synaptically occurs in the axonal presynaptic-like structure of cultured CGCs, in which presynaptic proteins are accumulated^21,22^. In the cultured *Nrxn* TKO CGCs, the action potential-induced Ca^2+^ influx into the axon was impaired (Fig. 7). Therefore, it is likely that the defect of axonal BDNF-pHluorin release in *Nrxn* TKO CGCs was caused by the defect of actionpotential-induced Ca^2+^ influx into the axonal presynaptic-like structure (Fig. 7). Thus, these results suggest that NRXNs organize the Ca^2+^ influx machinery that is essential for the release of neurotrophic factors. Our results also provide a molecular insight into the organization and function of the neurotrophic factor release site. Previous studies of α-NRXN TKO, conditional NRXN TKO, or β-NRXN TKO mice showed that both α-NRXNs and β-NRXNs play a role in controlling presynaptic Ca^2+^ channels in several types of synapses via different mechanisms^4,7,8,47^. Here, the action-potential-induced Ca^2+^ influx into the axonal presynaptic-like structure was impaired in *Nrxn* TKO CGCs. Furthermore, this defect was rescued by the expression of NRXN1β(+S4) (Fig. 7). Conversely, β-NRXNs reportedly function at the synapse, where they regulate Ca^2+^ influx by controlling the tonic postsynaptic endocannabinoid signaling mediated by 2-AG^4^. Thus, the regulatory mechanism of axonal Ca^2+^ influx mediated by NRXNs in cultured CGCs appears to be distinct from the synaptic functions of NRXNs.

The synapse formation process has been studied well in the neuromuscular junction. During development, prior to innervation, acetylcholine receptors are clustered in the prospective synaptic region of muscle fibers, which is called muscle pre-patterning. In mammals, pre-patterning requires the muscle-specific receptor tyrosine kinase MuSK and the low-density lipoprotein receptor-related protein 4 (Lrp4). Interestingly, muscle Lrp4 also binds to motor axons and sends a retrograde signal to the motor nerve for presynaptic differentiation^48^. In the cerebellum, presynaptic elements of the PF that exhibit vesicular profiles are intrinsically formed in the absence of specific postsynaptic compartments^10,11^. This is reproducible in *in vitro* CGCs cultures, in which axonal presynaptic-like structures containing synaptic proteins are intrinsically formed without contacting postsynaptic structures^21–23^. A previous study suggested that the presynaptic-like structures containing synaptic vesicle clusters observed in the cultured CGCs may represent the early stages of the presynaptic structure, which provides local platforms for synapse formation, as the axonal structural changes induced by GluD2 preferentially occurred in these regions^49^. In cultured *Nrxn* TKO CGCs, the density and intensity of Bassoon, synapsin, and VGluT1 puncta along the axons were severely decreased, suggesting a crucial role for NRXNs in the formation of the presynaptic-like structure (Fig. 8). These defects, as well as other defects, were rescued by NRXN1β(+S4), but not by its C-terminal mutant (Figs 6–8). Thus, it is reasonable to assume that the primary function of NRXNs in CGCs is the regulation of the organization of the axonal presynaptic-like structure through their C-terminal region. NRXNs interact with CASK and Mint through their C-terminal region^2^. However, the signal transduction pathway underlying this phenomenon has not been elucidated. Therefore, further studies are required to address this issue. We previously found that the number and intensity of Bassoon and VGluT1 in the axonal varicosities are increased after the application of the recombinant tetrameric GluD2 N-terminal domain in wild-type CGCs, but not in Cbln1 KO CGCs, suggesting that presynaptic differentiation is induced by the NRXN–Cbln1–GluD2 interaction^23^. It was also suggested that presynaptic differentiation is induced by the multimerization of NRXNs via their postsynaptic ligands, such as GluD2–Cbln1 or NLGN1^23,50^. In contrast, multimerization of NRXNs does not seem to be necessary for the formation of the presynaptic-like structure, as NRXNs in the axons of CGCs lack contact with the postsynaptic ligands that are required for NRXN multimerization.

In conclusion, by analyzing conditional *Nrxn* TKO CGCs both *in vivo* and *in vitro*, we found novel function of NRXNs in CGCs survival. Our results suggest that, in CGCs, NRXNs play essential roles in the formation and function of presynaptic-like structures containing presynaptic proteins, independent of binding to their postsynaptic ligands. Moreover, our results suggest that the survival of CGCs is maintained by autocrine neurotrophic factors with a secretory machinery that is organized by NRXNs.

## Acknowledgements

We thank Dr. T. Furuichi for providing pEFBOS-BDNF-4×pHluorin, Dr. R. Sprengel for providing the plasmid containing iCre, Dr. H. Niwa for providing pCAGGS, Ms. R. Koike and Ms. A. Maeda for technical assistance. This work was supported by Takeda Science Foundation to T.U., grants from JSPS/MEXT KAKENHI (JP25282242 and JP19H03334 to T.U., JP16H04676 to M.M., and JP25282242 to K.T., JST CREST (JPMJCR12M5) to T.U. and S.F., and JST PRESTO to T.Y. and K.T.

## Author contributions

T.U. and K.S. constructed plasmid vectors. T.U., K.S., E. M., and T. K. performed cell culture experiments. T.U., M.Y., M. A., and K. S. generated *Nrxn1^flox/flox^; Nrxn2^flox/flox^; Nrxn3^flox/flox^* mice. M. A., and K. S. prepared *GluN2C^+/iCre^* mice. W.M. prepared anti-pan-Nrxn, anti-VGluT1, anti-VGluT2, and anti-calbindin antibodies. T.U. and K.S. performed histological, immunohistochemical experiments. T.U. performed behavioral and electron microscopy experiments. P.F. and M.A. performed β-galactosidase assay. T.U. wrote the paper with editing M.Y., T.Y., S.F., M.M., and K.T. T.U. and K.T. designed and supervised the study.

## Additional information

Correspondence and materials should be addressed to T.U. or K.T.

## Methods

### Generation of CGC-specific *Nrxn1, 2, 3* TKO mice

Bacterial artificial chromosome (BAC) clones prepared from the C57BL/6 strain carrying the last exon of the *Nrxn1, Nrxn2*, and *Nrxn3* genes (RP24-119B4, RP24-174J7, and RP24-273A7, respectively) were purchased from BACPAC Resources Center and used to construct gene-targeting vectors. Genetargeting vectors were constructed using the Quick and Easy BAC Modification kit (Gene Bridges) and the Multisite Gateway system (Invitrogen). The 1.8 kb loxP-frt-Neo-frt cassette fragment carrying the loxP sequence and the phosphoglycerate kinase 1 (*Pgk-1*) promoter-driven neomycin phosphotransferase (*neo*) gene flanked by two Flp recognition target (frt) sites^17^ was inserted 0.7 kb upstream of the last exon of *Nrxn1*. The loxP sequence was inserted immediately after the stop codon of *Nrxn1*. The targeting vector pTV-Nrxn1 contained the last exon of *Nrxn1* flanked by loxP sequences, a *neo* cassette flanked by frt sequences, the 4.2 kb upstream and 5.6 kb downstream genomic sequences, and the 4.3 kb pMC1DTpA^51^. The 1.8 kb loxP-frt-Neo-frt cassette fragment was inserted 0.5 kb upstream of the last exon of *Nrxn2*. The loxP sequence was inserted immediately after the stop codon of *Nrxn2*. The targeting vector pTV-Nrxn2 contained the last exon of *Nrxn2* flanked by loxP sequences, a *neo* cassette flanked by frt sequences, the 4.4 kb upstream and the 6.7 kb downstream genomic sequences, and the 4.3 kb pMC1DTpA. The 1.8 kb loxP-frt-Neo-frt cassette fragment was inserted 0.8 kb upstream of the last exon of *Nrxn3*. The loxP sequence was inserted immediately after the stop codon of *Nrxn3*. The targeting vector pTV-Nrxn3 contained the last exon of *Nrxn3* flanked by loxP sequences, a *neo* cassette flanked by frt sequences, the 5.6 kb upstream and the 5.7 kb downstream genomic sequences, and the 4.3 kb pMC1DTpA. The linearized targeting vectors were introduced into the embryonic stem (ES) cell line RENKA, which is derived from the C57BL/6 strain, and G-418-resistant clones were selected^52^. The generation of chimeric mice was performed as described previously^52^. Briefly, the recombinant ES cells were injected into 8-cell-stage embryos of the CD-1 mouse strain. The embryos were cultured to blastocysts and transferred into the uterus of the pseudopregnant CD-1 mice. The resulting mice were mated to B6-Tg(CAG-FLPe)36 mice of the C57BL/6 strain^53^, to eliminate the *neo* gene from the genome through Flp/frt-mediated excision. The resulting *Nrxn1^+/flox^, Nrxn2^+/flox^*, and *Nrxn3^+/flox^* mice were crossed with each other to obtain *Nrxn1^flox/flox^; Nrxn2^flox/flox^; Nrxn3^flox/flox^* mice. To generate CGC-specific *Nrxn* knockout mice, *Nrxn1^flox/flox^; Nrxn2^flox/flox^; Nrxn3^flox/flox^* mice were crossed with E3CreN mice (*GluN2C^+/iCre^*), in which the codon-improved Cre recombinase (iCre) was specifically expressed in CGCs^17^. The resulting GluN2C^*+/iCre*^; *Nrxn1^+/flox^; Nrxn2^+/flox^; Nrxn3^+/flox^* mice were further crossed with *Nrxn1^flox/flox^; Nrxn2^flox/flox^; Nrxn3^flox/flox^* mice to yield *GluN2C^+/iCre^; Nrxn1^flox/+^*; *Nrxn2^flox/flox^*; *Nrxn3^flox/flox^* and *GluN2C^+/+^; Nrxn1^flox/flox^; Nrxn2^flox/flox^; Nrxn3^flox/flox^* mice. Male *GluN2C^+/iCre^; Nrxn1^+/flox^; Nrxn2^flox/flox^*; *Nrxn3^flox/flox^* mice were crossed with female *GluN2C^+/+^; Nrxn1^flox/flox^; Nrxn2^flox/flox^; Nrxn3^flox/flox^* mice, and the resulting mice were used in the experiments, unless otherwise specified. The *Nrxn1^+/flox^, Nrxn2^+/flox^*, and *Nrxn3^+/flox^* alleles were identified by PCR using the following primers: 5’–TGCCATGTACAAGTACAGGAAC–3’ and 5’–TTTATGAATGCGTGCGGTCCTC–3’, 5’–GTGCTATTTGCGCCCTCCGCC–3’ and 5’–TGTGAGCACGTGCCCTGCCTGAC–3’, and 5’–TGTTCCGTAATGTGCCCACAGC–3’ and 5’–CAACCTTGCAGCATGGCTCACC–3’, respectively. The amplified DNA fragments were as follows: *Nrxn1^+^*, 540 bp; *Nrxn1^flox^*, 642 bp; *Nrxn2^+^*, 760 bp, *Nrxn2^flox^*, 862 bp; *Nrxn3^+^*, 524 bp; and *Nrxn3^flox^*, 626 bp. The *GluN2C^iCre^* allele was identified by PCR using primers 5’–TAGGAAGGTTCTCTGTACTCTGGC–3’, 5’–CACAGATTTGGGTGAGGATGCTGC–3’, and 5’–GACTTGGTCAAAGTCAGTGCGTTC–3’. The amplified DNA fragments were as follows: *GluN2C^+^*, 362 bp; and *GluN2C^iCre^*, 570 bp. PCR was performed using the following conditions. *Nrxn1* and *GluN2C^iCre^*: one cycle of 94°C for 2 min, followed by 35 cycles of 98°C for 10 s, 60°C for 10 s, and 68°C for 35 s; *Nrxn2* and *Nrxn3*: one cycle of 94°C for 2 min, followed by 30 cycles of 98°C for 10 s and 68°C for 30 s. All PCRs were performed using KOD FX Neo polymerase (TOYOBO). The animal care and experimental protocols were reviewed by the Committee for Animal Experiments and approved by the president of Shinshu University (Authorization No. 280017 and 290072) and the Dean of the School of Medicine, University of Tokyo (Authorization No. 9), and conducted in accordance with the Guidelines for the Care and Use of Laboratory Animals of Shinshu University and University of Tokyo.

### Western blot analysis

Cerebellar and forebrain homogenates were prepared from 8-week-old mice. The brains were homogenized with homogenate buffer containing 0.32 M sucrose, 1 mM NaHCO_3_, 1 mM MgCl_2_, 0.5 mM CaCl_2_, and protease inhibitors using a Teflon-glass homogenizer, followed by centrifugation at 710 × *g* for 10 min at 4°C, as described previously^54^. Proteins were quantified using the Quick Start Bradford Protein Assay (Bio-Rad). Equal amounts of protein were separated by SDS–PAGE, transferred to a PVDF membrane, and incubated with rabbit anti-pan NRXN (0.42 μg/ml)^55^ and rabbit anti-β-actin (Abcam) (1:2000) antibodies, followed by incubation with a horseradish-peroxidase-conjugated secondary antibody, respectively. Immunoreactive proteins were visualized using the ECL Select Western Blotting Detection System (GE Healthcare). Signals were detected on a Las-4000 mini instrument (GE Healthcare) and quantified using the ImageQuant TL image analysis software (GE Healthcare).

### Histology and immunohistochemistry

Under deep anesthesia, mice were fixed transcardially with 4% paraformaldehyde (PFA) in phosphate-buffered saline, pH 7.4. Parasagittal cerebellar sections (50 μm in thickness) were prepared for immunohistochemistry using a microslicer (VT1200S; Leica Biosystems). The sections were stained with goat anti-VGluT1 (0.5 μg/ml)^56^, guinea pig anti-VGluT2 (1 μg/ml)^57^, and rabbit anti-calbindin (1 μg/ml) antibodies, followed by incubation with species-specific donkey Alexa Fluor 488-conjugated, donkey Alexa Fluor 647-conjugated (Thermo Fisher Scientific), and donkey Cy3-conjugated (Jackson ImmunoResearch) antibodies. The sections were counterstained with 1 μg/ml 4’, 6-diamidino-2-phenylindole (DAPI) (SIGMA) for 5 min. Fluorescence images of the whole cerebellum were acquired by a BZ-X700 microscope (Keyence). Fluorescence images of the cerebellar cortex were acquired using a confocal laser-scanning microscope (TCS SP8, Leica Microsystems) with an HC PL APO 40×/1.10 NA water-immersion lens (Leica Microsystems). Paraffin sections (5 μm thick) were prepared using a sliding microtome (REM-710, YAMATO) and stained with hematoxylin. The images were acquired using a microscope (DMLB 11888011, Leica Microsystems) equipped with a CCD camera (DFC 450C; Leica Microsystems).

### TUNEL assay

Terminal deoxynucleotidyl transferase-mediated dUTP nick end labeling (TUNEL) histochemistry was performed using the ApopTag Peroxidase In Situ Apoptosis Detection Kit (Merck Millipore), according to the instructions of the manufacturer. Briefly, 5 μm paraffin sections were incubated in PBS containing proteinase K at 22°C for 15 min, then treated with 3% H_2_O_2_ for 5 min, followed by incubation with reaction buffer containing DIG-labeled dUTP and terminal deoxynucleotidyl transferase at 37°C for 1 h. After the TUNEL reaction was terminated, the sections were incubated with a peroxidase-conjugated anti-DIG antibody, followed by incubation with peroxidase substrate buffer containing the DAB substrate and counterstaining with 0.5% methyl green. The images were acquired using a microscope (DMLB 11888011) equipped with a CCD camera (DFC 450C).

### β-galactosidase assay

The E3CreN mouse line (*GluN2C^+/iCre^*) was crossed with the Cre-inducible lacZ reporter mouse line (CAG-CAT-Z11)^58^. Brain slices prepared from the offspring carrying both the iCre and CAG-CAT-Z11 genes (*GluN2C^+/iCre^*; CAG-CAT-Z11/+) were stained for β-galactosidase activity as described previously^59^.

### Electron microscopy

Under deep pentobarbital anesthesia, mice were perfused transcardially with 2% PFA/2% glutaraldehyde in 0.1 M phosphate buffer, pH 7.4. The cerebellum was then removed and cut sagittally into 300-μm-thick sections using a microslicer (VT1200S). The sections were postfixed with 1% osmium tetroxide in 0.1 M phosphate buffer for 1 h at 4°C, followed by incubation with 1% uranyl acetate for 40 min at room temperature. After dehydration, the sections were embedded in Epon 812 (Nisshin EM). Parasagittal ultrathin sections (70 nm thick) were prepared using an ultramicrotome (Ultracut N ultramicrotome, Leica Microsystems) and stained with uranyl acetate for 5 min and Reynold’s lead citrate solution for 3 min. Electron micrographs were taken by a JEM-1400 electron microscope (JEOL).

### Antibody production

The rabbit anti-synapsin I polyclonal antibody was produced by the SIGMA custom antibody production services. The sequence of the antigenic polypeptide was NYLRRRLSDSNFMANLPNGYMTDLQRPQP, corresponding to the amino-terminus of synapsin I^60^. The specificity of the antibody was confirmed by western blotting using a mouse brain homogenate (Supplementary Figure 8).

### Construction of expression vectors

The coding sequence of the double human influenza hemagglutinin (2×HA) epitope tag (YPYDVPDYAYPYDVPDYA) was inserted immediately after the signal sequence of NRXN1β(+S4) and NRXN1β(−S4) by PCR with the following primers: 5’–GGATCCGGTACCGCCACCATGTACCAGAGGATGCT–3’, 5’–GGATCCTCGAGTCAGACGTAATACTCCTTATC–3’, 5’–CCAGATTATGCTTACCCCTACGACGTGCCCGATTATGCCTCCAGTTTGGGAGCGCAC–3’, and 5’–GTCGTAGGGGTAAGCATAATCTGGAACATCAT–3’ using pCAG-HA-NRXN1β^12^ and pCAG-HA-NRXN1β(−S4)^12^ as templates, and cloned into the *Kpn*I-*Xho*I sites of pCAG-1, to yield pCAG-2HA-NRXN1β(+S4) and pCAG-2HA-NRXN1β(−S4), respectively.

The DNA fragment carrying the extracellular and transmembrane regions of NRXN1β(+S4) was amplified by PCR with the primers 5’-GGATCCGGTACCGCCACCATGTACCAGAGGATGCT-3’ and 5’-TTATCCCGCCGTTTCATGGCATAGAGGAGGATGAG-3’ using pCAG-2HA-NRXN1β(+S4) as a template. The DNA fragment carrying the C-terminal cytoplasmic region of N-cadherin was amplified by PCR with primers 5’-CTATGCCATGAAACGGCGGGATAAAGAGCGC-3’ and 5’-GGATCCTCGAGTCAGACGTAATACTCCTTATC-3’ using pN-cadherin^61^ as a template. The amplified fragments were joined by overlap extension PCR and cloned into the *Kpn*I-*Xho*I sites of pCAG-1, to yield pCAG-2HA-NRXN1β(+S4)/N-cad. The DNA fragment carrying the extracellular region of NRXN1β(+S4) was amplified by PCR with primers 5’–ATCATTTTGGCAAAGCCACCATGTACCAGAGGATG–3’ and 5’–NRXN-GPI-R, CACAAAGTGAGCTGCGCCAGTGGTACTGCTGGACT–3’ using pCAG-2HA-NRXN1β(+S4) as a template. The GPI-anchored sequence of NCAM120 was amplified by RT-PCR with primers 5’–CAGTACCACTGGCGCAGCTCACTTTGTGTTCAGGAC–3’ and 5’–CCTGAGGAGTGAATTTCAGAGCAGAAGAAGAGTCA–3’ using mRNA prepared from an ICR mouse as a template. The amplified fragments were joined by overlap extension PCR and cloned into the *Eco*RI-*Xho*I sites of pCAG-1 using the GeneArt Seamless Cloning and Assembly Kit (Thermo Fisher Scientific), to yield pCAG-2HA-NRXN1β(+S4)-GPI. The DNA fragment carrying the extracellular region of NRXN1β(+S4) was amplified by PCR with primers 5’–ATCATTTTGGCAAAGCCACCATGTACCAGAGGATG–3’ and 5’–CCTTATAGCCAGTGGTACTGC–3’ using pCAG-2HA-NRXN1β(+S4) as a template. The DNA fragment carrying the transmembrane and cytoplasmic regions of IL-2Ra was amplified by RT-PCR with primers 5’–GCAGTACCACTGGCTATAAGGTAG–3’ and 5’–TGAGGAGTGAATTGCCTAGATGGTTCTTCTGCTC–3’ using mRNA prepared from an ICR mouse as a template. The amplified fragments were joined by overlap extension PCR and cloned into the *Eco*RI-*Xho*I sites of pCAG-1 using the GeneArt Seamless Cloning and Assembly Kit, to yield pCAG-2HA-NRXN1β/IL-2Ra. The entire coding sequence of GCaMP6s was amplified by PCR with primers 5’–GGATCCGCCACCATGGGTTCTCATCATCATCA–3’ and 5’–GTAGCCGAACCGGTCTTCGCTGTCATCATTTGTAC–3’ using pGP-CMV-GCaMP6s (Addgene plasmid # 40753; http://n2t.net/addgene:40753; RRID:Addgene_40753) as a template. The DNA fragment carrying rat VAMP2 (aa 2-116) was amplified by PCR with primers 5’–CAGCGAAGACCGGTTCGGCTACCGCTGCCACCGTC–3’ and 5’–GAATTCTTAAGTGCTGAAGTAAACG–3’ using pCAG-EGFP-VAMP2^12^ as a template. The amplified fragments were joined by overlap extension PCR and cloned into pCAG-1, to yield pCAG-GCaMP6s-VAMP2. The coding sequence of iCre^62^ was cloned into *Kpn*I-*Bam*HI sites of pCAG-1^12^, to yield pCAG-iCre. The entire coding sequence of iCre was amplified by PCR with primers 5’–ACCCACTGCCCCTTGGATCCGCCACCATGGTGCCCA–3’ and 5’–ATAAGCTTGATATCGGAATTCTCAGTCCCCATCCT–3’ using pCAG-iCre as a template, and cloned into the BamHI-EcoRI sites of the pFSy(1.1)GW lentiviral vector (Addgene plasmid # 27232; http://n2t.net/addgene:27232; RRID:Addgene_27232), to yield pFSy-iCre. The 9.0 kb BamHI-EcoRI fragment from pFSy(1.1)GW was blunt-ended with the Klenow fragment of DNA polymerase and ligated to yield an empty pFSy lentiviral vector. The entire coding sequences of 2HA-NRXN1β(+S4), 2HA-NRXN1β(−S4), 2HA-NRXN1β(+S4)-GPI, 2HA-NXN1β/IL-2Ra, 2HA-NRXN1β(+S4)/N-cad, and NRXN1α(+S4) were amplified by PCR using pCAG-2HA-NRXN1β(+S4), pCAG-2HA-NRXN1β(−S4), pCAG-2HA-NRXN1β(+S4)-GPI, pCAG-2HA-NRXN1β/IL-2Ra, pCAG-2HA-NRXN1β(+S4)/N-cad, and pNRXN1α-V5^63^ as templates, followed by cloning into the BamHI-EcoRI sites of pFSy(1.1)GW using the GeneArt Seamless Cloning and Assembly Kit or restriction enzymes, to yield pFSy-HA-NRXN1β(+S4), pFSy-2HA-NRXN1β(−S4), pFSy-2HA-NRXN1β(+S4)-GPI, pFSy-2HA-NRXN1β/IL-2Ra, pFSy-2HA-NRXN1β(+S4)/N-cad, and pFSy-NRXN1α(+S4), respectively. The sequences of the primers used in this experiment were as follows: 2HA-NRXN1β(+S4), 5’–ACCCACTGCCCCTTGCCACCATGTACCAGAGGATG–3’ and 5’–ATAAGCTTGATATCGTCAGACGTAATACTCCTTAT–3’; 2HA-NRXN1 β(−S4), 5’–GGATCCGGTACCGCCACCATGTACCAGAGGATGCT–3’ and 5’–CCGCTCGAGTCAGACGTAATACTCCTTATCCTTG–3’; 2HA-NRXN1β(+S4)-GPI, 5’–ACCCACTGCCCCTTGCCACCATGTACCAGAGGATG–3’ and 5’–ATAAGCTTGATATCGTCAGAGCAGAAGAAGAGTC–3’; 2HA-NRXN1 β(+S4)/N-cad, 5’–ACCCACTGCCCCTTGCCACCATGTACCAGAGGATG–3’ and 5’–ATAAGCTTGATATCGTCAGTCGTCACCACCGCCG–3’; 2HA-NRXN1 β/IL-2Ra, 5’–ACCCACTGCCCCTTGCCACCATGTACCAGAGGATG–3’ and 5’–ATAAGCTTGATATCGCTAGATGGTTCTTCTGCTC–3’. The cDNA encoding NRXN1α lacking S4 [NRXN1α(−S4)] was constructed via PCR-based mutagenesis using pFSy-NRXN1α(+S4) as a template.

### Lentivirus production

Lentiviruses were produced by transfection of Lenti-X 293T cells (Takara) with a lentiviral plasmid containing EGFP, iCre, or NRXN cDNAs, and the helper plasmids pMDLg/pRRE (Addgene plasmid # 12251; http://n2t.net/addgene: 12251; RRID: Addgene_12251), pRSV-Rev (Addgene plasmid # 12253; http://n2t.net/addgene: 12253; RRID: Addgene_12253), and pMD2.G (Addgene plasmid # 12259; http://n2t.net/addgene: 12259; RRID: Addgene_12259). Transfection was performed using Polyethylenimine (PEI) Max (Polysciences). Six micrograms of lentiviral plasmid and 2 μg of each helper plasmid were suspended in 0.6 mL Dulbecco’s Modified Eagle’s Medium (SIGMA) containing 36 μg of PEI Max for 15 min at room temperature. The DNA/PEI complexes were applied to Lenti-X 293T cells plated on a 10 cm diameter dish. Forty-eight hours after transfection, media were corrected and filtrated. Viruses were concentrated using Lenti-X concentrator (Takara) and resuspended with phosphate-buffered saline, pH 7.4. Viruses were stored at −80°C in aliquots until use.

### Cell culture and immunocytochemistry

Primary cerebellar granule cell (CGC) cultures were prepared from *Nrxn1^flox/flox^; Nrxn2^flox/flox^; Nrxn3^flox/flox^* or wild-type C57BL/6 mice at postnatal day 7 (P7), as described previously^64^. Briefly, the cells were cultured in Neurobasal-A (Thermo Fisher Scientific) supplemented with 2% B-27 (Thermo Fisher Scientific), 5% fetal calf serum (FCS), 100 U/mL of penicillin, 100 μg/mL of streptomycin, and 2 mM GlutaMax I (Thermo Fisher Scientific) for 24 h, and then cultured in the same medium without FCS. Cells were plated on cover slips coated with 30 μg/mL of poly-L-lysine and 10 μg/mL of mouse laminin at a density of 3 × 10^5^ cells or 2.5 × 10^4^ cells in a 24-well dish. The cultured CGCs were infected with either an empty lentivirus or a lentivirus expressing iCre at DIV1. The infected cells were fixed at DIV7 and immunostained with rabbit anti-pan NRXNs (1.6 μg/ml)^55^ and goat anti-tau (1:1000) (Santa Cruz Biotechnology) antibodies, followed by incubation with donkey Alexa Fluor 555-conjugated antigoat and donkey Alexa Fluor 488-conjugated anti-rabbit antibodies (Thermo Fisher Scientific). For double infection, the cultured CGCs were first infected with either the empty lentivirus or the lentivirus expressing iCre for 3 h at DIV1. After the first infections, the culture media were replaced with fresh media, followed by infecting with an empty lentivirus and a lentivirus expressing NRXNs or mutant NRXNs. For treatment with neurotrophic factors, 20 ng/ml of human BDNF (SIGMA), human NT-3 (Peprotech), human FGF acidic/FGF-1 (Peprotech), human FGF basic/FGF-2 (Peprotech), or human IGF-I (Cell Signaling) was applied to the cultured CGCs at DIV2. The cells were fixed with 4% PFA and 4% sucrose at DIV4, 7, or 10, and immunostained with a mouse anti-NeuN antibody (Merck Millipore). Fluorescence images were acquired with a confocal laser-scanning microscope (TCS SP8) using an HC PL APO CS2 20×/0.75 NA multiple-immersion lens (Leica Microsystems). The CGCs were identified based on NeuN signals and their morphological features, i.e., small size (5-10 μm in diameter) and regular round or ovoid shape^65^. For single-axon analysis, cationic liposome lipofectamine 2000 (Thermo Fisher Scientific) was used for sparse transfection. The cultured CGCs were transfected with pCAG-iCre and pCAG-TagRFP^66^, alone or together with pFSy-2HA-Nrxn1β(+S4) or p pFSy-2Nrxn1β(+S4)/N-cadC at DIV3. The cells were fixed at DIV7 and immunostained with mouse anti-Bassoon (1:500) (Stressgen), rabbit anti-synapsin I (1:1000), or rabbit anti-VGluT1 (0.37 μg/ml) antibodies^57^, followed by incubation with donkey Alexa Fluor 555-conjugated anti-mouse or anti-rabbit IgG antibodies (Thermo Fisher Scientific). Fluorescence images were acquired with a confocal laserscanning microscope (TCS SP8) using an HC PL APO CS2 20×/0.75 NA multiple-immersion lens (Leica Microsystems) or an APO CS2 100×/1.40 NA oil-immersion lens (Leica Microsystems) under constant conditions of laser power, iris, gain, z-steps, and zoom setting throughout the experiments. Images were collected from at least two independent experiments and all quantitative measurements were performed using the ImageJ 1.44v software. For the quantification of Bassoon, synapsin, and VGluT1 puncta, z-series optical sections of isolated axons were projected using the brightest point method and smoothened using the rank filter method. Bassoon, VGluT1, and synapsin puncta were defined as areas in which the staining signal intensity was 7, 4, and 4 times stronger than that of the background signals on the same field, respectively. Contiguous puncta were separated from each other using the “segmented particles” tool. For analysis of the localization of GCaMP6s-VAMP2 in cultured CGCs, these cells were transfected with TagRFP and GCaMP6s-VAMP2 at DIV3 using lipofectamine 2000. The cells were fixed at DIV7 and immunostained with a mouse anti-Bassoon antibody, followed by incubation with donkey Alexa Fluor 555-conjugated anti-mouse and rabbit Alexa Fluor 488-conjugated anti-GFP (Thermo Fisher Scientific) antibodies. Fluorescence images were acquired with a confocal laser-scanning microscope (TCS SP8) using an APO CS2 100×/1.40 NA oil-immersion lens. High-magnification images were deconvolved using Huygens Essential version (Scientific Volume Imaging).

### Behavioral test

A rotarod apparatus consisting of a rod with a diameter of 3.2 cm (RRAC-3002; O’Hara & Co., Tokyo, Japan) was used to measure motor coordination. Four-week-old male mice were placed on the rod, which was rotated at 15 rpm, and the latency to fall (retention time) was measured with a cutoff time of 5 min. Mice were trained for 3 consecutive days, receiving 3 trials per day with an intertrial interval of 2 h.

### Time-lapse imaging of BDNF secretion

For time-lapse imaging experiments, the CGCs prepared from *Nrxn1^flox/flox^; Nrxn2^flox/flox^; Nrxn3^flox/flox^* pups at P7 were plated on a glass-bottom culture dish coated with 30 μg/mL of poly-L-lysine and 10 μg/mL of mouse laminin at a density of 1.6 × 10^5^ cells/cm^2^. The cultured CGCs were transfected with pEFBOS-BDNF-4×pHluorin^67^ and pCAG-TagRFP, together with pCAG-iCre or pCAGGS^68^, at DIV3 using lipofectamine 2000. Four or five days after transfection, time-lapse imaging was performed on a confocal laser-scanning microscope (TCS SP8) using an APO CS2 100×/1.40 NA oil-immersion lens equipped with an auto-focusing stage (Leica Microsystems) and a warm chamber (Tokai Hit) under constant conditions of laser power, iris, gain, z-steps, and zoom setting throughout the experiments. Before imaging, media were with HEPES-buffered saline solution containing 140 mM NaCl, 5.4 mM KCl, 5 mM CaCl_2_, 0.8 mM MgCl_2_, 10 mM glucose, and 10 mM HEPES, pH 7.4. Depolarization-evoked BDNF secretion was induced by a HEPES-buffered saline solution containing 45 mM NaCl, 100 mM KCl, 5 mM CaCl_2_, 0.8 mM MgCl_2_, 10 mM glucose, and 20 mM HEPES, pH 7.4. All quantitative measurements were performed using the ImageJ 1.44v software. The images were collected sequentially at 30 s intervals. For the quantification of BDNF-pHluorin puncta along the axons of CGCs, z-series optical sections were projected using the brightest point method and smoothened using the uniform filter method. BDNF-pHluorin puncta were defined as areas in which the signal intensity was 4 times stronger than that of background signals on the same axon. Contiguous puncta were separated from each other using the “segmented particles” tool. Average fluorescence events before (2 min recording, 4 events) and after (12 min recording, 24 events) stimulation were quantified.

### Ca^2+^ imaging

For Ca^2+^ imaging experiments, the CGCs were plated on a glass-bottom culture dish as described above. The cultured CGCs prepared from *Nrxn1^flox/flox^; Nrxn2^flox/flox^; Nrxn3^flox/flox^* P7 pups were transfected with pCAG-iCre, pCAG-GCaMP6s-VAMP2, and pCAG-TagRFP with or without pFSy-2HA-NRXN1β(+S4) or pFSy-2HA-NRXN1β(+S4)/N-cad at DIV3 using lipofectamine 2000. Imaging was performed at DIV7 or 8 on a laser-canning confocal microscope (LSM7 Live, Carl Zeiss) using a 63×/1.4 NA plan apochromat objective lens (Carl Zeiss). TagRFP fluorescence was excited with a mercury lamp with a filter set, and CGCs and their axons were morphologically identified based on TagRFP fluorescence signals. GCaMP6s-VAMP2 fluorescence was excited by a 488 nm laser, and the collecting emission passed through a high-pass filter of 495 nm. The pinhole was adjusted to 1 A.U. GCaMP6s-VAMP2 fluorescence images were acquired at a rate of 15 Hz. All experiments were performed at room temperature and neurons were stimulated by field stimulation using nichrome wire in HEPES-buffered saline solution containing 140 mM NaCl, 5.4 mM KCl, 2 mM CaCl_2_, 1 mM MgCl_2_, 10 mM glucose, and 10 mM HEPES, pH 7.4. Stimulus trains were applied at 2, 5, 10, 17, 25, 50, and 75 stimuli at a rate of 20 Hz. To confirm the action-potential-stimulated Ca^2+^ influx, the cultures were treated with 0.5 μM tetrodotoxin (Tocris). The time course of fluorescence transients was obtained from regions of interest, as defined by manual selection of GCaMP6s-VAMP2 fluorescence using ZEN2010 (Carl Zeiss). Changes in GCaMP6s-VAMP2 fluorescence images were quantified as the mean region of interest and expressed as % ΔF/F, where F is the baseline fluorescence level and ΔF is the change in fluorescence intensity from this value.

### Statistical analysis

The results of at least two independent experiments were subjected to statistical analyses. No statistical method was used to determine sample size. Statistical significance was evaluated using the Kruskal–Wallis test followed by the post-hoc Steel-Dwass test, two-way or one-way ANOVA followed by Tukey’s or Dunnett’s post hoc test, or Student’s *t*-test using the R software (R Core Team, 2017). Statistical significance was assumed when *P* < 0.05.

### Data availability

The data generated in this experiment are available from the corresponding author upon reasonable request.

