## Supplementary Figures for "Neurexins play a crucial role in cerebellar granule cell survival by organizing autocrine machinery for neurotrophins"

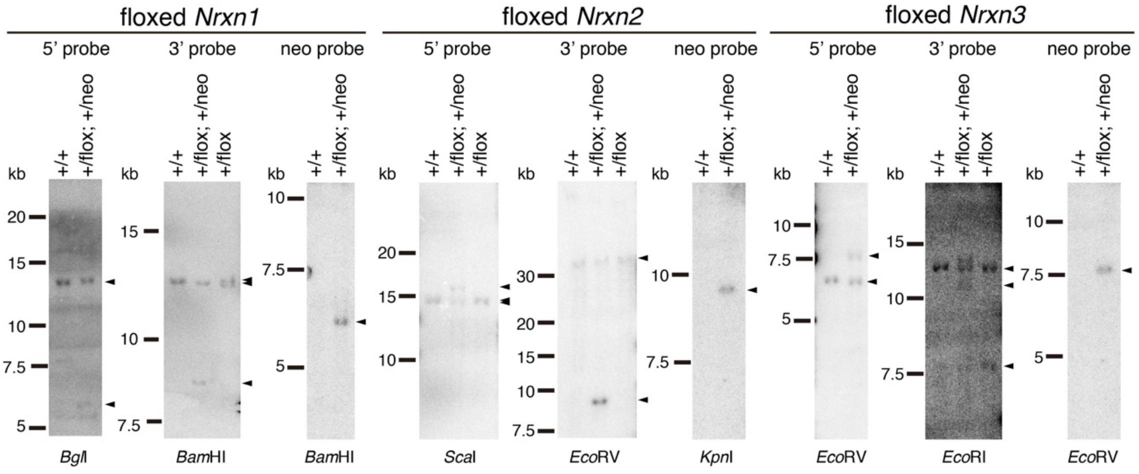


**Supplementary Figure 1. Southern blot analysis of genomic DNA.**

Genomic DNAs from $Nrxn1^{+/+}$, $Nrxn1^{+/flox;\ +/neo}$, $Nrxn1^{+/flox}$, $Nrxn2^{+/+}$, $Nrxn2^{+/flox;\ +/neo}$, $Nrxn2^{+/flox}$, $Nrxn3^{+/+}$, $Nrxn3^{+/flox;\ +/neo}$, and $Nrxn3^{+/flox}$ mice were digested with indicated restriction enzymes and analyzed by a southern blotting using 5', 3', and neo probes, respectively.

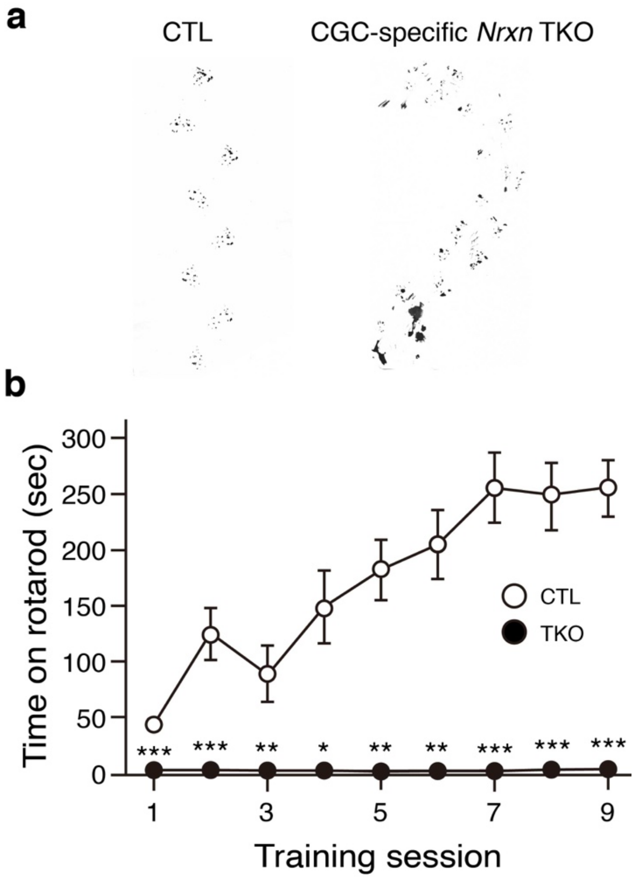


**Supplementary Figure 2. Footprint patterns and performance in the constant-speed rotarod test of CGC-specific *Nrxn* TKO mice**

**a**, Representative footprint patterns from control and CGC-specific *Nrxn* TKO mice. **b**, The constant-speed rotarod test. The time that mice remained on the rotarod before falling was measured as a function of trial session. The open and closed circles represent control (n = 6) and CGC-specific *Nrxns* TKO (n = 6) mice, respectively. All values represent the mean ± s.e.m. ***$P < 0.001$, **$P < 0.01$, *$P < 0.05$; two-way repeated ANOVA (training session × genotype interaction, $F$ (8, 107) = 20.5, $P = 2.5 \times 10^{-17}$), followed by *post hoc* two-tailed Student's *t*-test. CTL, control mice; TKO, CGC-specific *Nrxn* TKO mice.

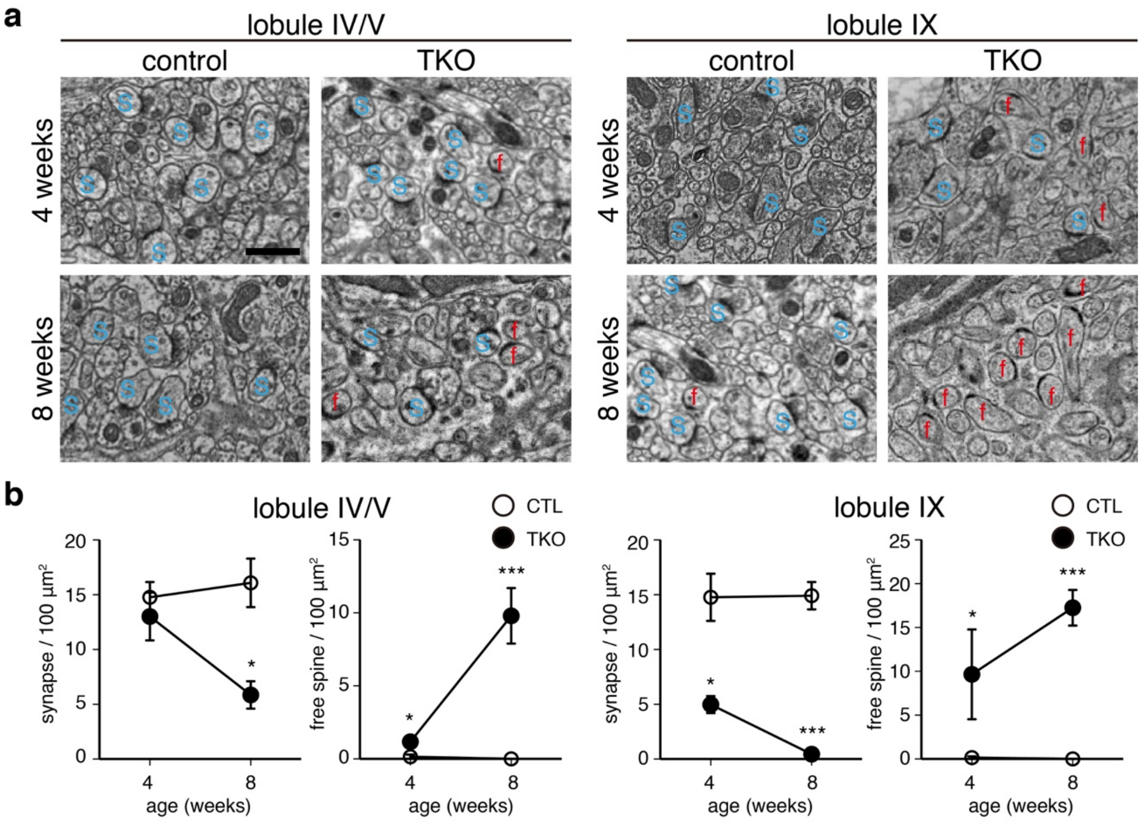


**Supplementary Figure 3. Structure of PF–PC synapses in CGC-specific *Nrxn* TKO mice**

**a**, Electron micrographs of the structure of PF–PC synapses of lobules IV/V and IX in 4-week-old or 8-week-old control and CGC-specific *Nrxn* TKO mice. S, synapse possessing spines that contact with the presynaptic terminal; f, free spines possessing PSD lacked presynaptic counterparts. Scale bar, 1 µm. **b**, Number of synapses and free spines in **a**. All values represent the mean ± s.e.m. ***$P < 0.001$, **$P < 0.01$, *$P < 0.05$; n = 3 each from three animals; two-way ANOVA (synapse in lobules IV/V, age × genotype interaction, $F$ (1, 11) = 5.5, $P$ = 0.048; free spine in lobules IV/V, age × genotype interaction, $F$ (1, 11) = 20.7, $P$ = 0.0019; synapse in lobule IX, genotype effect, $F$ (1, 11) = 80.8, $P = 1.9 \times 10^{-5}$; free spine in lobule IX, genotype effect, $F$

(1, 11) = 23.6, $P = 1.3 \times 10^{-3}$), followed by *post hoc* two-tailed Student's *t*-test (synapse, CTL vs. TKO at 4 and 8 weeks in lobule IV, $P$ = 0.54 and 0.016, respectively; free spine, CTL vs. TKO at 4 and 8 weeks in lobule IV, $P$ = 0.035 and 0.0067, respectively; synapse, CTL vs. TKO at 4 and 8 weeks in lobule IX, $P$ = 0.013 and $5.4 \times 10^{-4}$, respectively; free spine, CTL vs. TKO at 4 and 8 weeks in lobule IX, $P$ = 0.037 and 0.0011, respectively). CTL, control mice; TKO, CGC-specific *Nrxn* TKO mice.

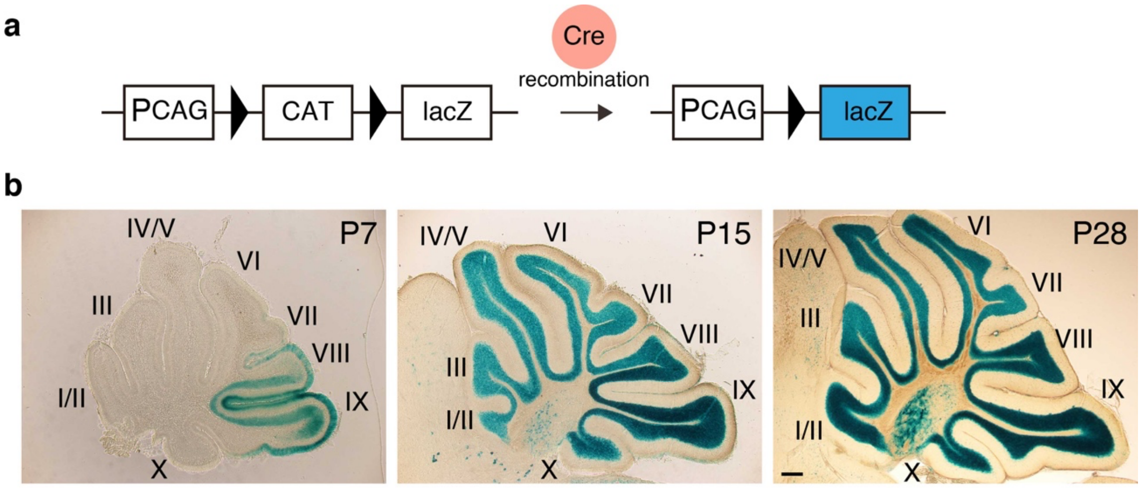


**Supplementary Figure 4. Cre recombinase activity in cerebellar granule cells in *GluN2C*$^{+/iCre}$ mice**

**a**, Schema of the Cre recombinase assay in vivo. Cre mediates recombination between two loxP target sites flanking the *CAT* gene, leading to the expression of the β-galactosidase gene (*lacZ*) driven by the CAG promoter (PCAG). **b**, Developmental time course of Cre activity in the cerebellum in *GluN2C*$^{+/iCre}$ mice. In *GluN2C*$^{+/iCre}$ mice, Cre-mediated recombination was first observed in lobules VIII and IX at postnatal day 7 (P7), and gradually expanded to other lobules as the cerebellum developed. Scale bar, 250 μm.

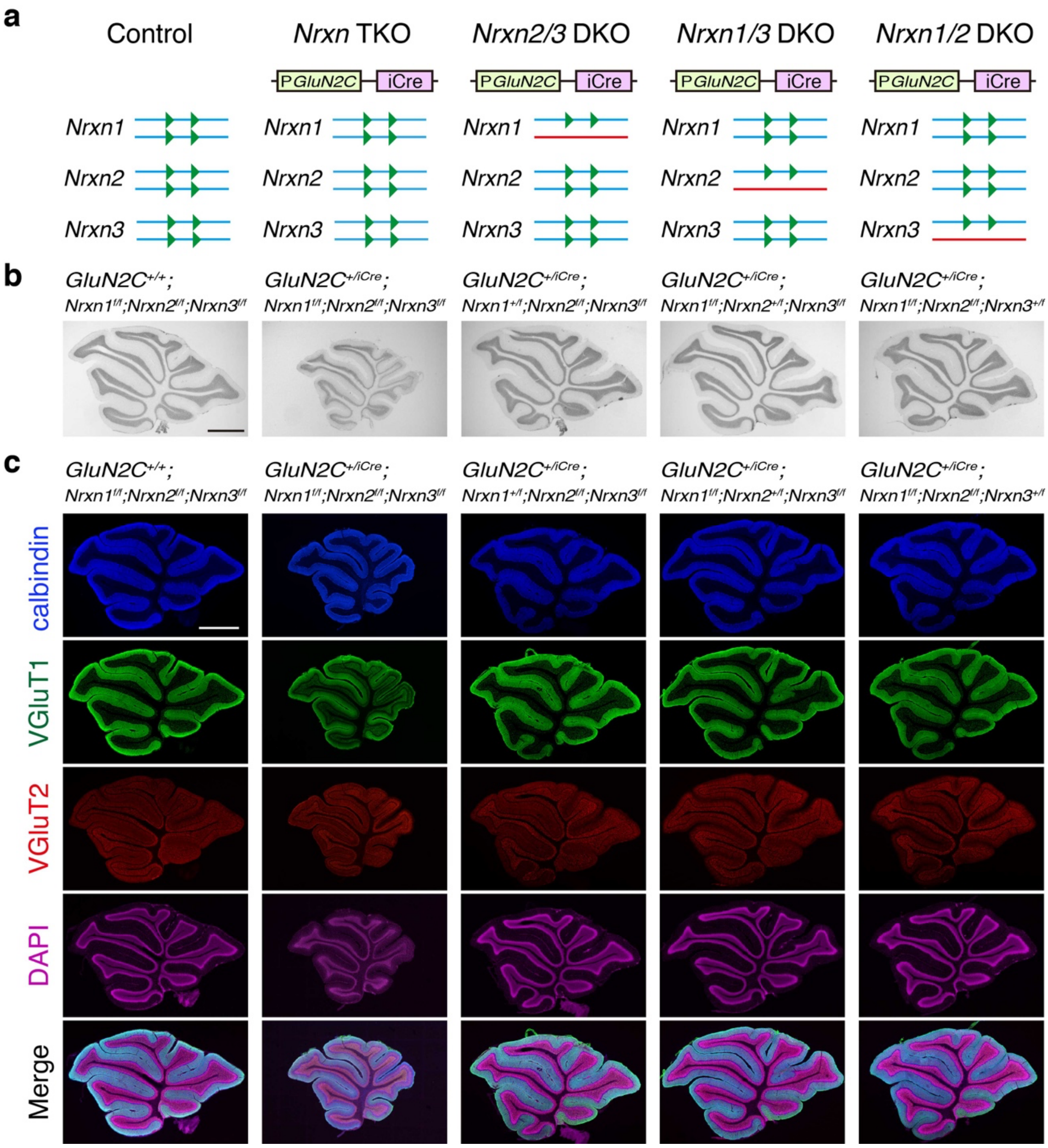


**Supplementary Figure 5. *Nrxn* DKO mice show apparently normal cerebellar structures**

**a**, Schematic representation of the *Nrxn1*, *Nrxn2*, and *Nrxn3* genes in *Nrxn* DKO mice. Floxed alleles are shown in blue lines containing *loxP* sequences (green triangles). Wild-type alleles are shown in red lines. Cre-mediated recombination leads to the deletion of *Nrxn* genes in floxed alleles. **b**, Histological cerebellar features of *Nrxn* DKO mice. Parasagittal cerebellar sections from 4-week-old control and CGC-specific *Nrxn* double DKO mice were stained with hematoxylin. **c**, Immunohistochemical analysis of *Nrxn* d DKO mice. Parasagittal cerebellar sections from 4-week-old control and CGC-specific *Nrxn* double DKO mice were stained with antibodies against VGluT1 (green), VGluT2 (red), Calbindin (blue), and DAPI (purple). The images were merged on the bottom. Scale bars, 1 mm in **b**,**c**.

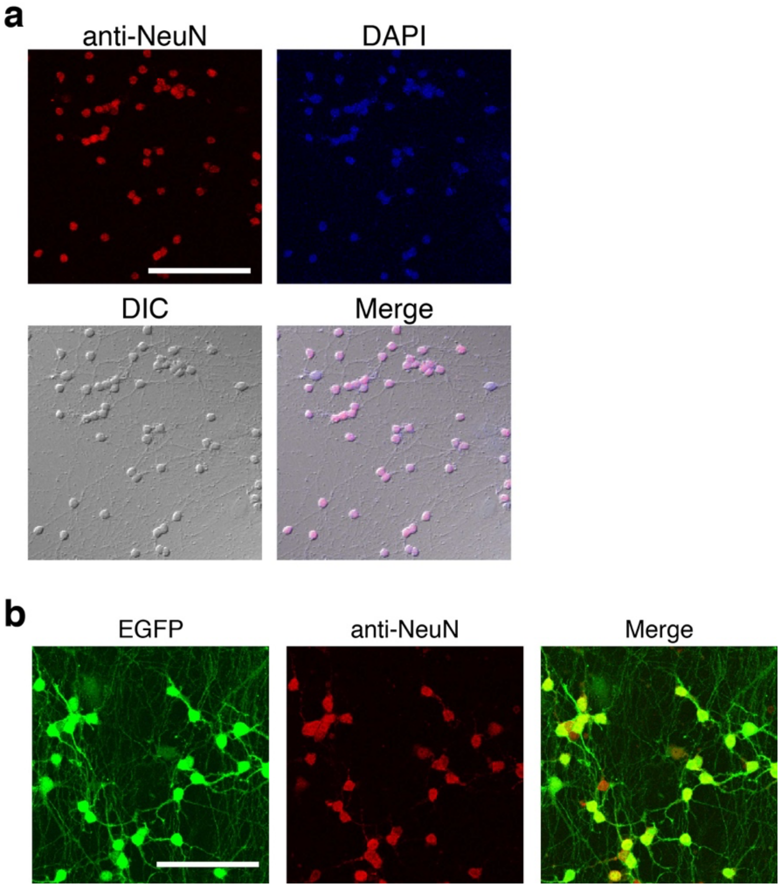


**Supplementary Figure 6. Population of cultured CGCs and lentivirus infection efficiency**

**a**, The cultured CGCs were prepared from P7 pups. The cells were stained with an antibody against NeuN. The CGCs were identified based on NeuN signals and their morphological features, i.e., small size (5−10 μm in diameter) and a regular round or ovoid shape. In our culture conditions, 95.9% ± 0.86% cells are CGCs (n = 3). **b**, Lentivirus infection efficiency in cultured CGCs. Cultured CGCs were infected with lentivirus-EGFP and immunostained with an antibody against NeuN. Infection efficiency (95.0% ± 0.37%, n = 3) was estimated by counting the percentage of EGFP-positive cells in NeuN-positive cells. Scale bars, 100 μm in **a** and 10 μm in **b.**

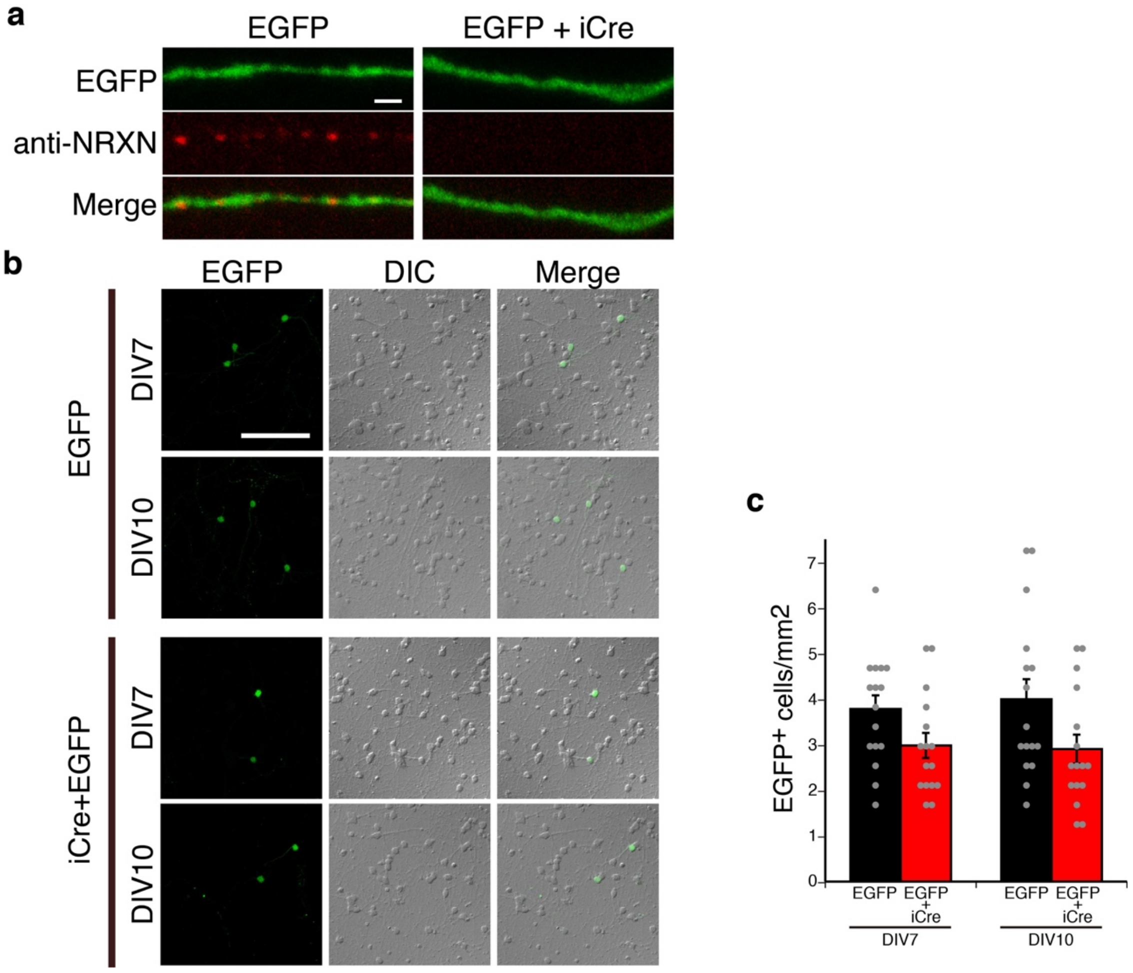


**Supplementary Figure 7. *Nrxn* TKO in a small population of cultured CGCs**

**a,** *Nrxn* TKO in sparsely transfected CGCs. The cultured *fNrxn* CGCs were sparsely transfected with EGPF alone or together with iCre. The transfected cells were immunostained with an antibody against NRXNs. The images of the axons of EGFP-positive CGCs are shown.

**b**, Effect of *Nrxn* TKO in a small population of cultured CGCs on cell survival. The cultured *fNrxn* CGCs were sparsely transfected with EGPF alone or together with iCre. Transfection efficiency (approximately 1.5%) was estimated by counting the percentage of EGFP-positive cells in EGFP transfected cultured CGCs. Fluorescence images (left panels) and DIC images (middle panels) are merged on the right. **c**, Quantification of number of EGFP positive cells in **b**. All values represent the mean ± s.e.m. n = 16 areas from three cultures, respectively; two-tailed Student's *t*-test (EGFP vs. EGFP+iCre at DIV7, $P = 0.058$; EGFP vs. EGFP+iCre at DIV10, $P = 0.054$). Scale bars, 100 μm in **a** and 1 μm in **c**.

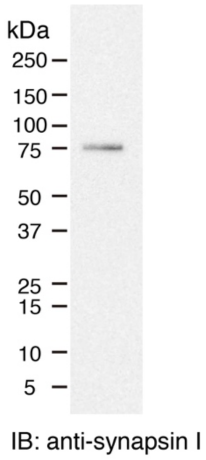


**Supplementary Figure 8. Specificity of the anti-synapsin I antibody**

A brain homogenate was separated by SDS–PAGE, transferred to a PVDF membrane, and incubated with the rabbit anti-synapsin I antibody, followed by incubation with a horseradish-peroxidase-conjugated secondary antibody. The antibody detected a band of ~75 kDa corresponding to the size of synapsin I[1,2].
